# Optimising synthetic cystic fibrosis sputum media for growth of non-typeable *Haemophilus influenzae*

**DOI:** 10.1101/2025.01.07.631681

**Authors:** Phoebe Do Carmo Silva, Darryl Hill, Freya Harrison

## Abstract

Non-typeable *Haemophilus influenzae* (NTHi) is an early pathogen isolated from the lungs of children with cystic fibrosis (CF). However, its role in the progression of CF lung infection is poorly understood. Additionally, whether it forms biofilms in the lungs of people with CF is an open question. The development of synthetic cystic fibrosis sputum media has given key insights into the microbiology of later CF pathogens, *Pseudomonas aeruginosa* and *Staphylococcus aureus*, through replicating the chemical composition of CF sputum. However, growth of NTHi in these media has not previously been reported. We show that NTHi grows poorly in three variants of synthetic cystic fibrosis sputum media commonly used to induce *in vivo* -like growth of *P. aeruginosa* and *S. aureus* (SCFM1, SCFM2 and SCFM3). The addition of NAD and hemin to SCFM1 and SCFM2 promoted the planktonic growth and biofilm formation of both laboratory and clinical NTHi isolates, and we were able to develop a modified variant of SCFM2 that allows culture of NTHis. We show that NTHi cannot be identified in an established *ex-vivo* model of CF infection, which uses SCFM and porcine bronchiolar tissue. This may in part be due to the presence of endogenous bacteria on the pig lung tissue which outcompete NTHi, but the lack of selective agar to isolate NTHi from endogenous bacteria, and the fact that NTHi is an exclusively human pathogen, make it hard to conclude that this is the case. Through spiking modified SCFM2 with filter sterilized lung homogenate, biofilm growth of clinical NTHi isolates was enhanced. Our results highlight that there are crucial components present in the lung tissue which NTHi require for growth, that are not present in any published variant of SCFM from the Palmer et al. 2007 lineage. Our results may inform future modifications to SCFM recipes to truly mimic the environment of CF lung sputum, and thus, to facilitate study of a wide range of CF pathogens.

**Data Summary:** The authors confirm that all supporting data, code and protocols have been provided within the article or through supplementary data files. All raw data has been uploaded to FigShare (https://doi.org/10.6084/m9.figshare.28175300.v1).

## Introduction

Cystic fibrosis (CF) is an autosomal genetic disorder, affecting over 70,000 individuals worldwide [1]. The accumulation of dehydrated mucus in the lungs harbors bacterial colonisation, resulting in chronic infection, a strong inflammatory response and lung damage [2]. The median life expectancy of a person with CF is currently 49 years in the United Kingdom [3], and many people with CF die prematurely from respiratory complications, despite treatment advances. The course of CF lung infection commonly begins with *Haemophilus influenzae* and *Staphylococcus aureus* in early childhood before progression to chronic infection with *Pseudomonas aeruginosa* during later childhood and adolescence [4].

There has been a dramatic increase in the survival of people with CF within the last decade thanks to a combination of antibiotic therapy, newborn screening and cystic fibrosis transmembrane conductance regulator (CFTR) modulators. CFTR modulators are now effective in correcting CFTR protein function in around 90% of people with CF [5][6][7]. There is increasing evidence that CFTR modulators reduce colonisation of later stage pathogens like *S. aureus* and *P. aeruginosa*[8]. However, whilst CFTR modulators decrease the prevalence of later stage pathogens, modulator therapy appears to result in an increase in prevalence of *H. influenzae* [9][10][11]. This means that investigating *H. influenzae* infection in CF lung mimicking conditions would be beneficial to better understand this changing landscape of CF lung infections. In this paper, we investigated the ability of *H. influenzae* to grow both planktonically and as a biofilm in a range of CF mimicking media to broaden the understanding of *H. influenzae* lung infection in people with CF.

Non-typeable *H. influenzae* (NTHi) are Gram-negative coccobacilli isolated from the respiratory tract in around 30% of children with CF [3][12]. Persistence of NTHi in the lower respiratory tract is commonly associated with periods of pulmonary exacerbation and chronic lung infection, contributing to the morbidity of people with CF [13][14] [15]. *H. influenzae* is defined by its requirement for Factors X (hemin) and V (nicotinamide adenine dinucleotide, NAD) to grow in laboratory conditions [16][17]. NTHi has emerged as the predominant group of *H. influenzae* strains in people with CF due to a lack of specific treatment, alongside a dramatic decline in invasive typeable *H. influenzae* disease due the development of a serotype b conjugate vaccine [18][19].

The existence of matrix-encased microbial communities known as biofilms is known to contribute to disease progression, pathogenesis and bacterial persistence in CF lung infections [20]. There is increasing evidence that *in vivo*, bacterial biofilms are increasingly tolerant to antibiotic therapy, surviving up to 100-1,000 fold higher antibiotic concentrations than planktonic cells [21]. Biofilms perpetuate chronic infection and recurrence of disease [22][23]. Mucin and extracellular deoxyribonucleic acid (eDNA) present in CF lungs have been demonstrated to provide a key bacterial niche for microcolony and biofilm formation *in vivo* [24][68]. Furthermore, hypoxic regions in the mucus have been demonstrated to increase transcription of efflux pumps, promote anaerobic respiration, reduce bacterial metabolism and induce biofilm formation in the well-studied CF pathogen, *P. aeruginosa* [22][25][26].

There is evidence that NTHi can form aggregates that are consistent with the appearance of a biofilm, but whether NTHi truly form biofilms, or if NTHi biofilms contribute to disease, is under debate [27][28][29][30]. There is a wealth of research into NTHi biofilm formation in otitis media and chronic obstructive pulmonary disease, whilst research on biofilm formation in CF is limited [31][32][33]. Biofilms containing NTHi have been directly visualised within the ear canal of children with otitis media using scanning electron microscopy [34], and shown to produce mucosal biofilms in a chinchilla model of otitis media using scanning electron microscopy [33] and confocal laser scanning microscopy [35]. Furthermore, NTHi has been shown to grow as a biofilm within the chronic obstructive pulmonary disease ferret lung model by confocal fluorescent imaging [36]. Conversely, little is known about *H. influenzae* in the context of CF colonisation: although there is evidence that suggests NTHi can form biofilm aggregates in the lungs [23][37], this does not confirm whether NTHi biofilms are stable within a CF lung environment. As such, we strived to gain an understanding of if *H. influenzae* form biofilms in a CF lung environment.

The nutritional environment of CF sputum is poorly replicated in standard laboratory growth media. To better recapitulate the CF lung environment *in vitro*, synthetic cystic fibrosis sputum media (SCFM) was developed based on the concentrations of ions, free amino acids, lactate and glucose found in CF sputum [38]. This SCFM, later termed SCFM1, has been well documented in the literature to be capable of supporting high density bacterial growth and biofilm formation of *P. aeruginosa* and *S. aureus in vitro*and in an *ex vivo* model of lung biofilm using porcine tissue [39] [40] [41] [42]. However, SCFM1 lacks several polymers and molecules present in CF sputum such as mucin and eDNA, which contribute to the viscosity of CF mucus and biofilm structures of pathogens within the CF lung. The presence of mucin has been shown to be crucial for microcolony development and bacterial biofilm formation [43], whilst eDNA is known to be present in large amounts in in CF lung mucus, and high concentrations have been shown to be associated with increased biofilm viscoelasticity and more robust biofilms [44] [45]. SCFM1 was therefore adapted to form SCFM2, with additions of bovine maxillary mucin, salmon sperm DNA, dioleoylphosphatidylcholine (DOPC) and N-acetyl glucosamine (GlcNAc) to make up for previous shortcomings [46]. To create a further version, SCFM3, final additions of p-aminobenzoic acid, NAD+, adenine, guanine, xanthine, and hypoxanthine were added [47]. Although these SCFMs have been optimised for usage with *P. aeruginosa* and *S. aureus*, there is no published research investigating the growth of NTHi in any variant of artificial sputum media.

In this paper, we investigated the growth profiles of NTHi in several variants of SCFM. We hypothesized that the fastidious nature of NTHi may mean NTHi may require supplementation of additional factors to support growth within SCFM, such as NAD and hemin. Therefore, either fetal bovine serum (FBS) or purified hemin and NAD were added. We then investigated the growth of NTHi in an established model of CF lung infection [48]. This combines pig bronchiolar and alveolar tissue with SCFM to mimic the structural and chemical environment of a human CF lung, and has previously supported the studies of key CF pathogens *P. aeruginosa* and *S. aureus* [41][40][42][49]. This paper is the first publication that attempts to use the *ex-vivo* pig lung model (EVPL) to investigate the growth of NTHi. In summary, our results demonstrate that while some simple modifications of SCFM formulations can support growth and biofilm formation of NTHi, growth is relatively poor. Further, we found that the EVPL model in its current form is not suitable for culture of NTHi. Further investigation is needed if we are to optimise CF-mimicking media and models for research into the pathogenicity and management of NTHi in the CF lungs.

## Methodology

### Bacterial strains

All *H. influenzae* were cultured on supplemented brain heart infusion (sBHI) agar (Millipore) at 37°C in 5% CO_2_. Following immersion in a water bath at 50°C, autoclaved agar was supplemented with NAD (2 *µ*g/mL) (Merck) and hemin (10 *µ*g/mL) (Sigma), mixed, and 20 mL was added to 90 mm polystyrene petri dishes. sBHI plates were stored at 4°C.

The *H. influenzae* strains used in this study are listed in Supplementary Table 1. The reference strains NCTC 11315, NCTC 04560, NCTC 12699 and NCTC 12975 were obtained from the National Collection of Type Cultures (www.culturecollections.org.uk). The streptomycin resistant NTHi375 and a GFP-tagged clinical otitis media isolate was derived from a Finnish pneumococcal vaccine study on children undergoing tympanocentesis in 1994-1995 [50] [51]. When cultured, NTHi-375 was plated on sBHI agar supplemented with 2*µ*g/mL streptomycin. All *Haemophilus influenzae* clinical CF isolates were obtained from patients of the SCILD cohort at the institute of infectious disease, University of Bern and are listed in Supplementary table 2 [96][97].

The fluorescent *Pseudomonas aeruginosa* PAO1 was derived from the Holloway, 1955 study [52] and subcultured on Luria-Bertani agar overnight at 37°C.

### Synthetic Cystic Fibrosis Sputum Media

Synthetic Cystic Fibrosis Sputum Media 1 (SCFM1) was made following the recipe by Palmer *et al* [38], minus the addition of glucose (following [41] (Supplementary Table 3)).

Synthetic Cystic Fibrosis Sputum Media 2 containing bovine maxillary mucin (SCFM2) was made following the recipe of Turner *et al*. [46]. The recipe was modified slightly by removing the phenol extraction of salmon sperm deoxyribonucleic acid (DNA) (Sigma). Instead, salmon sperm DNA was sterilised by ultraviolet (UV) light in 30 mg aliquots for 20 minutes. This was done to simplify the recipe by removing the time-consuming and high-hazard step of phenol extraction (Supplementary Table 4), and followed advice from one of the original authors (K Turner, personal communication).

Modified Synthetic Cystic Fibrosis Sputum Media 2 with sialic acid replacing bovine maxillary mucin (SCFM2.SA) was primarily prepared following the recipe of the Turner et al. [46]. Modifications included UV sterilisation of salmon sperm DNA as per SCFM2, and replacing bovine maxillary mucin with sialic acid (n-acetylneuraminic acid (Neu5Ac)) (Millipore). A stock concentration of 45 mg/mL Neu5Ac in dimethylsulfoxide (DMSO) was used. In 100 mL of SCFM2.SA, 1 mL of this stock was added for a final concentration of 450 *µ*g/ mL (Supplementary Table 4). The replacement of mucin with sialic acid is more affordable than bovine maxillary mucin and also removes the need for a 4-hour UV sterilisation step and the potential contamination of SCFM2 with impurities present in commercial mucin.

For experiments, each of SCFM1, SCFM2 or SCFM2.SA was used exactly as described above; or supplemented with 2% fetal bovine serum (Gibco); or supplemented with NAD (2 *µ*g/mL) and hemin (10 *µ*g/mL).

Synthetic Cystic Fibrosis Sputum 3 was made using the recipe by Neve *et al*., [47]. Modified SCFM2.SA was used as a base, with the additions of p-aminobenzoate (200 *µ*M) NAD (10 *µ*g/ml), adenine, guanine, xanthine and hypoxanthine (10 *µ*g/ml) (Supplementary Table 4).

### Growth of bacteria

All *H. influenzae* strains were maintained on sBHI plates at 37°C in 5% CO_2_. A colony of *H. influenzae* was grown overnight in BHI broth supplemented with NAD (2 *µ*g/mL) and hemin (10 *µ*g/mL) at 37°C in 5% CO_2_. Bacteria were washed by centrifuging 1 mL of preculture in microfuge tubes for 1 minute at 10,500 rpm, removing the supernatant, adding 1 mL phosphate buffered saline (PBS) and vortexing to resuspend cells. The centrifuge and PBS wash steps were then repeated. The desired volume of overnight culture to achieve OD_600_ 0.1 was then added to fresh microfuge tubes and washed with PBS as before. Cells were then resuspended in the SCFM media of choice.

### Growth curves

All strains and isolates of *H. influenzae* were grown overnight as stated and adjusted to a desired optical density of OD_600_ 0.1 in SCFM media of choice. 200 *µ*L of each suspension was added to a flat-bottom Corning 96 well polystyrene plate, with four replica wells inoculated per condition. Media-only wells were used as a negative control. The plate was then incubated in a Tecan Spark 10M multimode plate reader at 37°C, 5% CO_2_ with shaking every 20 minutes. OD_600_ (nm) was read every 20 minutes for 24 hours. Carrying capacity (K) and growth rate (r) parameters were then extracted using the *growthcurver* package in RStudio [53]. For plotting and analysis, OD_600_ values were blanked on the starting OD_600_ of each well.

To determine the colony forming units (CFU) per well, following 24 hours of growth, 100 *µ*L from each well was removed and serially diluted from 10^0^ to 10^−7^. 10 *µ*L from each dilution were then plated on sBHI plates.

The colony forming units were then counted following static incubation for 24 hours at 37°C in 5% CO_2_.

### Static biofilm assay

*H. influenzae* were grown overnight, washed and re-suspended in media of choice to OD_600_ 0.1 as before. 100 *µ*L of each suspension was added to a flat-bottom Corning 96 well polystyrene plate (Sigma). Plates were incubated statically at 37°C and 5% CO_2_ for 48 hours. Media was then slowly aspirated from wells to remove planktonic cells and added to a fresh flat-bottom 96 well plate, leaving behind the biofilm. 100 *µ*L of PBS was added to each biofilm well, and the base of well was scraped with a pipette tip. Plates were sealed with Parafilm and sonicated for 10 minutes in a sonicating water bath (Grant XUBAL XUBA1, Grant Instruments (Cambridge Ltd.)). Following sonication, absorbance of both this plate and the aspirated planktonic cell plate were measured at OD_600_ in a Tecan SPARK 10M multimode plate reader.

For crystal violet staining, following aspiration of media and planktonic cells, 100 *µ*L of crystal violet (0.1% in PBS) (Sigma) was added to each well and incubated at room temperature for 15 minutes. Crystal violet was aspirated and 100 *µ*L of cell culture water (HyClone) was added to each well, aspirated, and fresh cell culture water was added. 100 *µ*L of 95% ethanol was then added to each well and incubated at room temperature for 15 minutes. The bottom of the wells were scraped with pipette tip and absorbance of each well measured at 590 nm in a Tecan SPARK 10M multimode plate reader.

Relative biofilm formation was calculated by taking this value, and dividing it by the OD_600_ of the aspirated planktonic cells.

### Biofilm assay using EBBA Biolight 680

EBBA BioLight 680 (Ebba BioTech) was diluted 1:100 in HyClone water and then 1:10 in SCFM of choice. Bacteria of interest (*H. influenzae* NCTC 11315, NCTC 4560, NCTC 12699 and NCTC 12975) were grown overnight, washed, and re-suspended in SCFM of choice to OD_600_ 0.1 as before. 100 *µ*L was added to black polystyrene transparent flat bottomed 96 well plates (Greiner) and incubated for 48 hours in a Tecan SPARK 10M multimode plate reader at 37°C in 5% CO_2_ with shaking, and OD_600_ fluorescence (ex/em 540/670 nm) read every 20 minutes.

### *Ex vivo* pig lung model (EVPL) dissection and infection

The EVPL model was prepared as previously described [54] [41][40]. Two independent lungs were used and provided by Taylors Butchers (Coventry, UK) and Steve Quigley and Sons Butchers Ltd (Cubbington, UK). Following dissection, bronchiole or alveolar tissue pieces were placed in 400 *µ*L SCFM1 with ampicillin, or ampicilllin (50 *µ*g/mL) + gentamicin (50 *µ*g/mL) and incubated at 37°C for 24 hours on a rocking platform. These antibiotics were chosen based on preliminary sequencing of commensal bacteria present on the lung tissue (data not shown). Following incubation, the tissue was washed twice with PBS and once with SCFM. Following the wash stages, the tissue was transferred to a fresh transparent 24-well plate with an SCFM2.SA.XV agarose pad, and infected as previously described prior to addition of 500 µL of SCFM2.SA.XV and incubation for 48 hours at 37°C and 5% CO_2_.

### Bacterial recovery from EVPL

Tissue was homogenised by bead beating as previously described [55][40]. To determine bacterial load, lung homogenate was serially diluted in PBS, and plated on sBHI agar plates, which were then incubated for 24 hours at 37°C and 5% CO_2_ to determine CFU. Agar plates contained either sBHI, or sBHI containing vancomycin (5 *µ*g/mL), bacitracin (300 *µ*g/mL) and clindamycin (1 *µ*g/mL) (VBC).

### Visualisation of fluorescently-tagged bacteria in EVPL

The fluorescence of uninfected tissue, NTHi-375 and PAO1 infected tissue was measured in an 8 x 8 grid at 0, 24 and 48 hours post infection, at 484 ex / 505 em in a Tecan SPARK 10M multimode plate reader using a flat-bottom transparent Corning 24-well plate (Costar).

### Synthetic CF sputum lung homogenate media

Pig lung square sections were acquired through dissection and washed as per the EVPL model, above. Each tissue section was homogenised in 1 mL of SCFM2.SA media in a FastPrep-24 machine (MP Biomedicals, UK) for 40 s at 4 m/s. Lung homogenate was then transferred to a sterile 1.5 mL microfuge tube and centrifuged for 1 minute at 10,500 rpm to remove tissue debris. The supernatant was removed and then diluted 1:4 in SCFM and filter sterilised through a 3 mm 0.2 *µ*m syringe filter (Fisher Scientific, China) with a 2 ml syringe (Becton Dickinson, Spain) into a sterile 50 mL tube. NAD (2 *µ*g/mL) and hemin (10 *µ*g/mL) were then added. This was refrigerated for up to 1 month.

### Statistical analysis

All data was analysed using RStudio (Mac OS Ventura 13.5.2 + Version 1.4.1717) (RStudio, https://www.rstudio.com) using general linear models followed by Analysis of Variance (ANOVA) and either post-hoc Dunnett, Estimated Marginal Means for pairwise comparisons or Tukey Honestly Significant Difference (HSD) tests as specified in the text.

## Results

### Adding NAD and hemin to SCFM2.SA significantly increases growth rate and carrying capacity of NTHi laboratory strains

The growth profiles of four laboratory strains, *H. influenzae* NCTC 12699, NCTC 12975, NCTC 4560 and NCTC 11315, were monitored in a total of nine different SCFM variations (SCFM1, SCFM2 and SCFM2.SA) alone or supplemented with either 2% FBS or NAD and hemin) over 24 hours to determine which SCFM variant was most suitable for NTHi growth. Growth curves are available in Supplementary Figure 1.

It was previously suggested that NAD is present in CF sputum, at concentrations up to 15 *µ*M [47], whilst hemin has been determined as ‘potentially available’ [46]. Haemoptysis, which is blood streaking in sputum, is observed in 60% of CF patients, and often a marker of airway inflammation, pulmonary exacerbation and declining lung function, with up to 35 *µ*M of hemin potentially found in CF sputum [56] [57]. Serum, containing NAD, hemin and additional factors which may influence bacteria growth, is also likely to be present in CF sputum due to vascular leakage following tissue damage from airway inflammation [57], and increased levels of serum proteins have been detected in the sputum of people with CF compared to healthy individuals [58]. We thus compared formulations of SCFM containing either purified NAD and hemin alone, or serum.

In SCFM2.SA, mucin in the original SCFM2 recipe was replaced with N-Acetylneuraminic acid at 450 *µ*g/ml to increase affordability and ease of preparation. NTHi binds host mucin via sialic acid through its lipooligosaccharide to evade the immune response, and this sialic acid is the most common sialic acid found in human mucin [59] [60]. The concentration was determined based on reports of sialic acid plasma concentrations of 402-550 *µ*g/ml in CF patients [61].

When growth of NTHi was investigated in these SCFM variations, there was a significant difference in carrying capacity and growth rate between media variations for all strains. Carrying capacity of all four NTHi laboratory strains when grown in SCFM2.SA supplemented with NAD + hemin (SCFM2.SA.XV) was significantly higher than when grown in unmodified SCFM1 (**Figure 1**). There was some variation but, overall, SCFM2.SA resulted in a higher carrying capacity and growth rate compared to modified SCFM2 and unmodified SCFM1, and within SCFM2.SA, adding FBS or XV increased the carrying capacity. Meanwhile, the addition of FBS or XV to SCFM1 and SCFM2 did not enhance carrying capacity or growth media relative to these medias with no additions.

**Figure 1:**
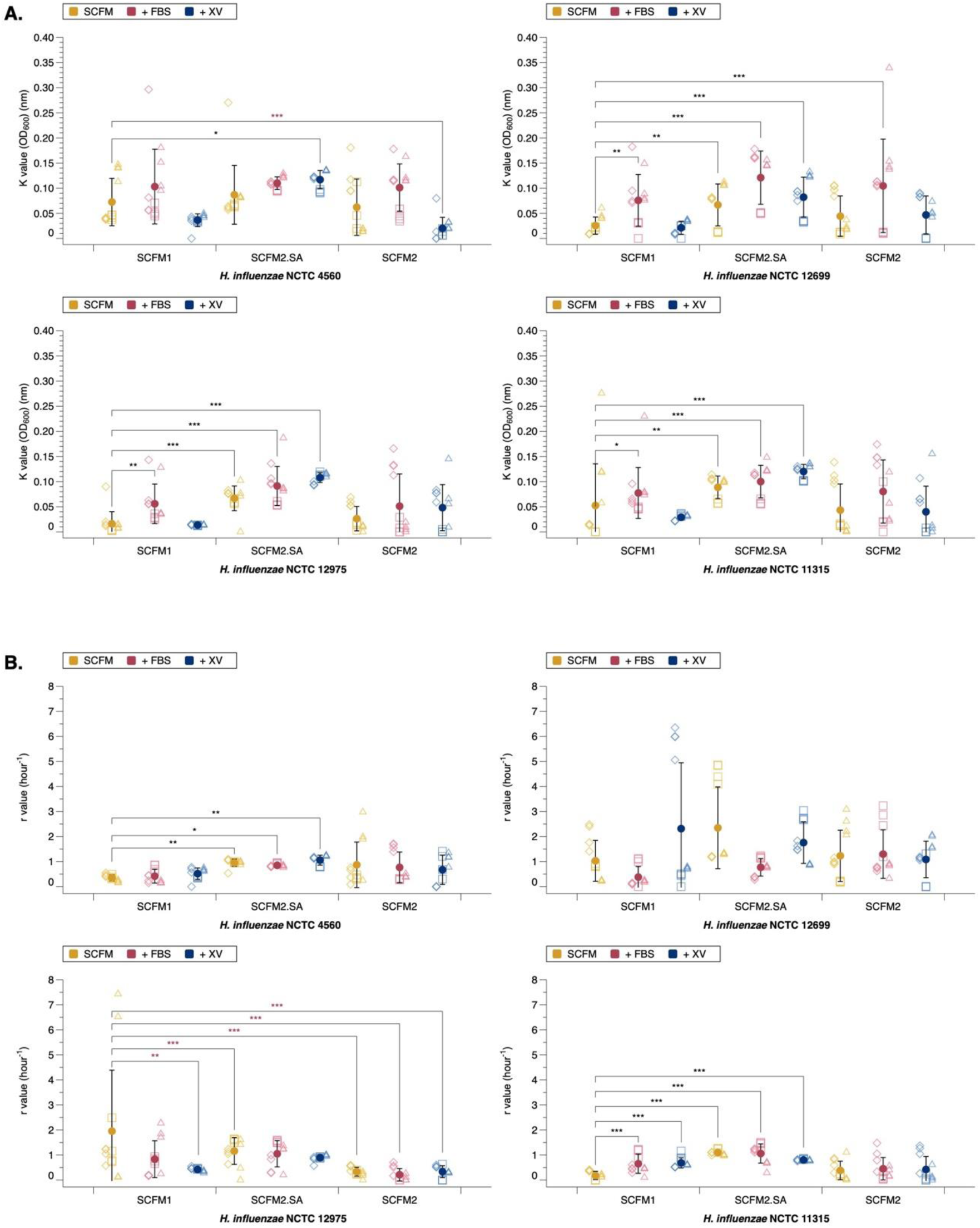
The influence of different SCFM media on the growth of four laboratory *H. influenzae* strains. Laboratory *H. influenzae* strains were grown in either SCFM1, SCFM2 or SCFM2.SA alone or supplemented with either fetal bovine serum or NAD and hemin. Carrying capacity and growth rate were calculated from 3 independent experiments (demonstrated by different shapes), with the mean and standard deviation shown by the solid circle. Significance was defined as *p* ≤ 0.05. An ANOVA was run to test the effect of strain type on carrying capacity (NCTC 12975 *F*_8,97_ = 13.17, *p* = < 0.001, NCTC 12699 *F*_8,97_ = 13.31, *p* = < 0.001, NCTC 11315 *F*_8,97_ = 8.70, *p* = < 0.001, NCTC 4560 *F*_8,97_ = 12.84, *p* = < 0.001). Post-hoc Dunnett analysis identified which media variations resulted in a significant change in K compared to SCFM1. A significant increase in K for all strains was identified when grown in SCFM2.SA.XV (NCTC 12975 *p* = < 0.001, NCTC 12699 *p* = < 0.001, NCTC 13315 *p* = < 0.001, NCTC 4560 *p* = 0.03). K was also significantly increased in all strains except NCTC 4560 when grown in SCFM2.SA (NCTC 12975 *p* = < 0.001, NCTC 12699 *p* = 0.007, NCTC 11315 *p* = 0.003), SCFM2.SA.FBS (all except NCTC 4560 *p* < 0.001) and SCFM1.FBS (NCTC 12975 *p* = 0.002, NCTC 12699 *p* = 0.002, NCTC 11315 *p* = 0.023). A significant increase in K was seen with NCTC 12699 when grown in SCFM2.FBS (*p* < 0.001) and a significant decrease in K was seen with NCTC 4560 when grown in SCFM2.XV) (*p* < 0.001). For growth rate, an ANOVA was run to test the effect of strain type on growth rate (NCTC 12975 *F*_8,97_ = 7.05, *p* = < 0.001, NCTC 12699 *F*_8,97_ = 3.87, *p* = < 0.001, NCTC 11315 *F*_8,96_ = 12.44, *p* = < 0.001, NCTC 4560 *F*_8,97_ = 4.14, *p* = < 0.001). Dunnett analysis identified which media variations resulted in a significant change in r compared to SCFM1. Growth rate was significantly increased when grown in SCFM2.SA (NCTC 4560 *p* < 0.001, NCTC 11315 *p* = 0.006), SCFM2.SA.FBS (NCTC 4560 *p* < 0.001, NCTC 11315 *p* = 0.022), SCFM2.SA.XV (NCTC 4560 *p* < 0.001, NCTC 11315 *p* = 0.001) SCFM1.FBS (NCTC 11315 *p* < 0.001) and SCFM1.XV (NCTC 11315 *p* < 0.001). Growth rate of NCTC 12975 was significantly reduced in all variants of SCFM2 (all *p* < 0.001) as well as SCFM1.XV (*p* = 0.003). There was no change in r for NCTC 12699 in any media variant.

These results suggest that the modified SCFM has a more prominent effect on enhancing carrying capacity than growth rate, in particularly SCFM2.SA.XV. SCFM2.SA.XV was the only media variant to significantly promote carrying capacity over SCFM1 in all four strains, and growth rate in two strains (NCTC 11315 and NCTC 4560). SCFM2.SA and SCFM2.SA.FBS increased growth rate in the same two strains and carrying capacity in all but NCTC 4560. On the contrary, the original SCFM2 recipe produced highly variable results and did not significantly enhance either carrying capacity or growth rate except in one case, with NCTC 12699 when supplemented with FBS. Supplementing SCFM1 with FBS enhanced carrying capacity in 3 strains (all but NCTC 4560) and growth rate in one strain (NCTC 11315).

NTHi is known to reach an average bacterial load of 10 to 10 CFU/ml when recovered from broncoalveolar lavage [62] [63] [64]. Therefore, it was investigated whether SCFM2.SA.XV could support a high bacterial load (CFU). Bacterial load of NTHi in SCFM2.SA.XV averaged 10 CFU/ml for NCTC 12975, 10 CFU/ml for NCTC 12699 and NCTC 4560, and 10 CFU/ml for NCTC 11315 (Supplementary Figure 2). This was slightly higher than in SCFM1, in which bacterial load was 10 CFU/ml for NCTC 12975, NCTC 12699 and NCTC 4560, whilst NCTC 11315 reached 10 CFU/ml. This suggests both SCFM1 and SCFM2.SA.XV can support NTHi growth to levels similar to that found in CF.

### Biofilm forming ability of NTHi *in vitro* is poor in SCFM, and fetal bovine serum promotes planktonic growth over biofilm growth

Several species of bacteria are known to form biofilms in the CF lung environment, and this often contributes to persistent lung infection [65]. We investigated whether NTHi are able to form biofilms in SCFM media *in vitro*, and to what extent its capability to form biofilm differs between the media variants. Biofilm formation was assessed by static growth in a flat-bottomed 96-well plate and staining with crystal violet at 48 hours; this is a high-throughput method to determine biomass based on optical density [66]. The relative biofilm allocation (amount of stain bound by biofilm / optical density of planktonic culture) was then calculated.

At 48 hours, there was measurable NTHi biofilm formation (Supplementary Figure 3). The relative biofilm allocation of each NTHi laboratory strain was determined in each SCFM variant to highlight whether any media variant promoted biofilm growth specifically, or promoted both planktonic and biofilm growth indiscriminately. A significant increase in relative biofilm, compared with unmodified SCFM1, was seen only seen in NCTC 12699 when grown in SCFM2.SA.XV. However, the addition of FBS, both to SCFM1 and SCFM2.SA significantly reduced relative biofilm formation for all NTHi strains (**Figure 2**). SCFM2.SA also significantly reduced relative biofilm formation of NCTC 11315 and NCTC 12699. Overall, this suggests that FBS promotes planktonic growth of bacteria over biofilm formation.

**Figure 2:**
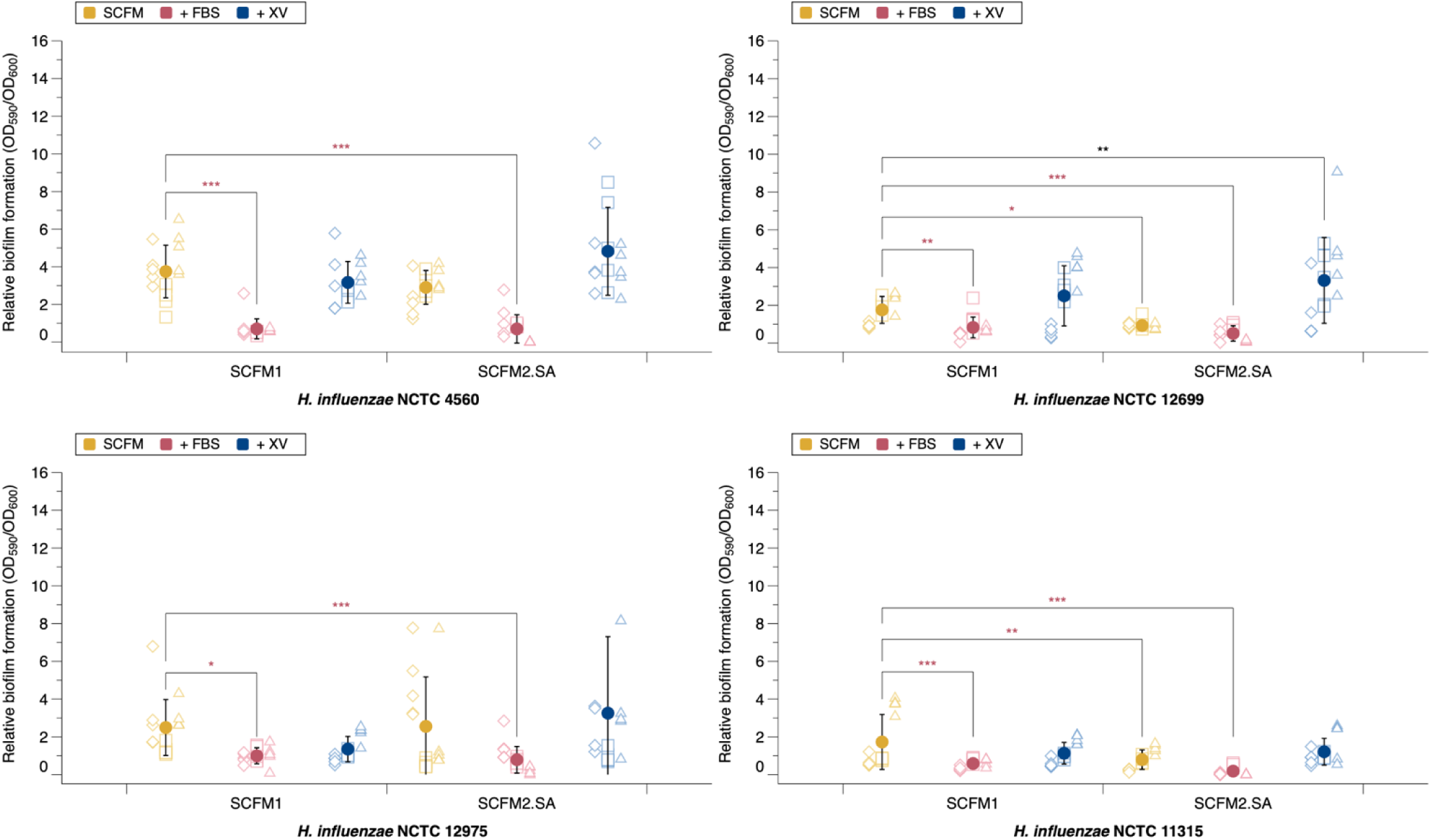
Relative biofilm allocation of four laboratory *H. influenzae* strains in varying SCFM conditions at 48 hours. Absorbance of biofilm was measured at 590 nm and planktonic at 600 nm. Mean and standard deviation was calculated from 3 independent experiments (demonstrated by different shapes), and shown by the solid circle. Significance was defined as *p* ≤ 0.05. An ANOVA was run to test the effect of media variation on relative biofilm formation for each strain (NCTC 12975 *F*_5,82_ = 5.93, *p* = *<*0.001, NCTC 12699 *F*_5,82_ = 20.21, *p* = *<*0.001, NCTC 11315 *F*_5,82_ = 21.41, *p* = *<*0.001, NCTC 4560 *F*_5,82_ = 37.51, *p* = *<*0.001). Post-hoc Dunnett analysis identified which media and strain comparisons resulted in a significant change in relative biofilm formation compared to SCFM1. There was a significant reduction in relative biofilm formation with SCFM1.FBS (NCTC 12975 *p <* 0.016, NCTC 12699 *p <* 0.005, NCTC 11315 *p <* 0.001, NCTC 4560 *p <* 0.001) and SCFM2.SA (all *p <* 0.001). SCFM2.SA.XV significantly promoted relative biofilm formation in NCTC 12699 (*p* = 0.007). SCFM2.SA significantly reduced relative biofilm formation of NCTC 11315 (*p* = 0.034) and NCTC 12699 (*p* = 0.002).

Overall, these results suggest that FBS may prevent biofilm formation of NTHi and may not be an ideal supplement for SCFM. As such, despite FBS previously promoting the *in vitro* proliferation of NTHi, the presence of FBS hinders biofilm growth. SCFM2.SA.XV was the only media variant to significantly promote biofilm formation, although in only one laboratory strain.

### Extracellular matrix production of *H. influenzae* is increased when grown in SCFM2.SA.XV

Crystal violet staining is a high-throughput and well-established assay used for quantifying biofilms *in vitro*. However, a downside to crystal violet is that it can stain bacterial cells and biofilm matrix, and it stains cells regardless of whether they are alive or dead [27]. As such, to specifically investigate biofilm matrix development, we utilised EBBA BioLight 680, a chemically modified optotracer which binds to the extracellular matrix of Gram negative bacteria and fluoresces only when bound [67]. NTHi was grown statically for 48 hours in SCFM2.SA and SCFM2.SA.XV with EBBA BioLight 680 to compare biofilm formation in these media. FBS was also tested as supplement, but was found to bind to EBBA BioLight (data not shown). Biofilm formation of NTHi was apparent when grown in SCFM2.SA and SCFM2.SA.XV. With all NTHi strains, fluorescence intensity of EBBA Biolight 680 increased when grown in SCFM2.SA.XV over SCFM2.SA.

In SCFM2.SA, fluorescence peaked around 8 hours at 7,000 RFU for NCTC 4560, 10 hours and 3,000 RFU for NCTC 11315 and less than 1,000 RFU for NCTC 12699 which did not appear to peak. On the other hand, biofilm formation in SCFM2.SA.XV was most apparent with NCTC 4560 and NCTC 11315. With NCTC 4560, fluorescence peaking at around 22,000 RFU at 16 hours before gradually decreasing up to 48 hours whilst NCTC 11315 peaked around 15,000 RFU at around 18 hours (**Figure 3**). NCTC 4560 and NCTC 11315 produced similar fluorescence profiles to each other, while fluorescence with NCTC 12699 and NCTC 12975 were similar to each other but much lower, peaking around 10,000 RFU at 16 hours and 5,000 RFU at 20 hours respectively. This suggested NCTC 12699 and NCTC 11315 do not produce as much extracellular matrix as NCTC 4560 or NCTC 11315, and may be weaker biofilm formers in SCFM. The time at which fluorescence peaked differed in all strains, but ranged between 16 to 20 hours. For both NCTC 4560 and NCTC 11315, fluorescence peaked before dropping entirely in some repeats. The peak of fluorescence appears later than when NTHi reached carrying capacity from the growth curves between 5 and 9 hours, consistent with the manufacturer’s claim that EBBA BioLight does not bind to the bacterial cell directly, and is indicative of biofilm formation by binding the extracellular matrix.

**Figure 3:**
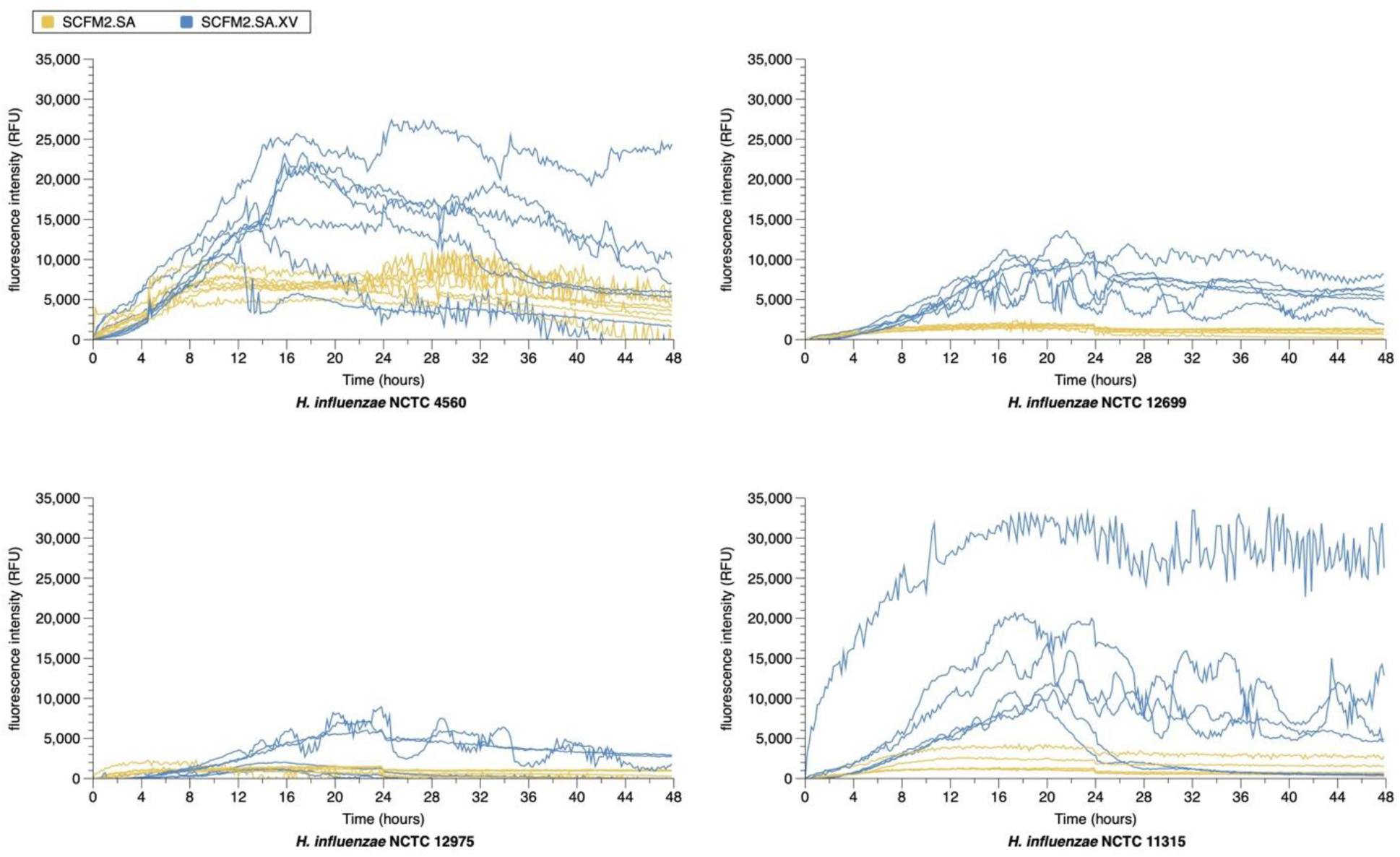
48-hour growth of *H. influenzae* with extracellular matrix marker EBBA BioLight. NTHi laboratory strains NCTC 4560, NCTC 12699, NCTC 12975 and NCTC 11315 were grown statically for 48 hours in SCFM2.SA and SCFM2.SA.XV. Fluorescence was measured at 540/570 ex/em (nm) over two independent experiments.

Overall, this data suggests all NTHi are capable of forming biofilms in SCFM2.SA and SCFM2.SA.XV, determined by production of an extracellular matrix. However, more extracellular matrix was produced in in SCFM2.SA.XV, with biofilm formation peaking between 16-20 hours. From this, SCFM2.SA.XV was determined to be the most suitable SCFM variant for use with NTHi for further *in vitro* work.

### Removing extracellular DNA from SCFM2.SA.XV significantly reduced growth of NTHi

Extracellular DNA is one of the major contributing factors to the progression of infection as well as biofilm formation in people with CF, and can be derived from both bacterial and host origins. The majority of eDNA in the lungs has been proposed to be due to neutrophil extracellular traps, but also comes from other sources such as dead host immune cells, lysed bacteria and bacterial DNA released through quorum sensing regulation [98]. Large amounts of eDNA are known to be present in CF lung mucus, and high concentrations have been shown to be associated with increased biofilm visco-elasticity and more robust biofilms [44] [45]. To determine the importance of eDNA in NTHi growth, the growth profiles of NTHi were investigated in SCFM2.SA.XV with and without eDNA. Growth curves are available in Supplementary Figure 4.

The removal of eDNA from SCFM2.SA.XV significantly reduced carrying capacity and growth rate in all laboratory strains (**Figure 4**). This suggests that eDNA is a crucial component to promote NTHi growth. The importance of eDNA in growth and biofilm has been previously suggested in the literature for other CF pathogens such as *P. aeruginosa* [68].

**Figure 4:**
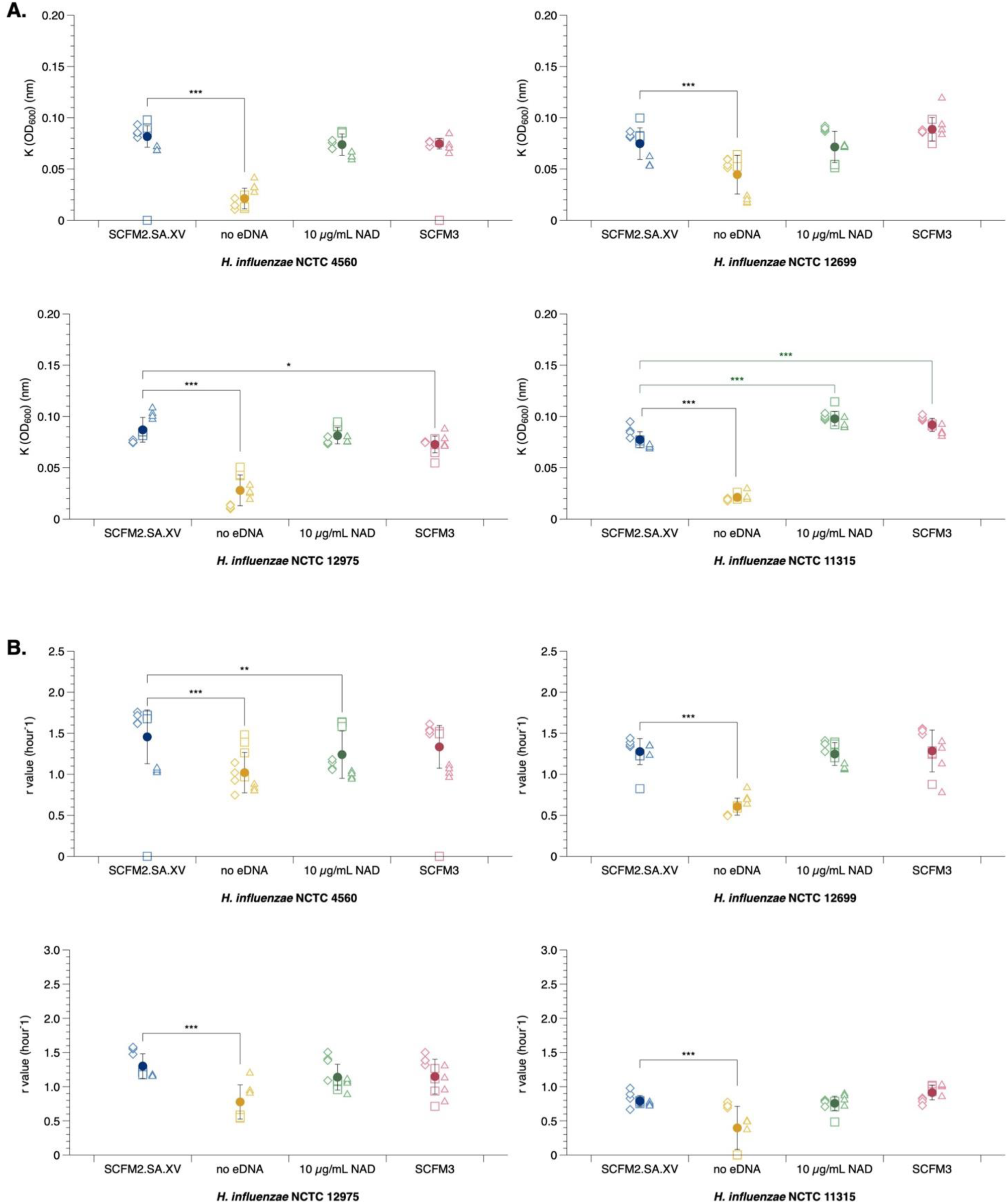
Carrying capacity and growth rate of *H. influenzae* in different SCFM medias. NTHi laboratory strains NCTC 4560, NCTC 12699, NCTC 12975 and NCTC 11315 were grown statically for 24 hours in either SCFM2.SA.XV, SCFM2.SA.XV minus eDNA, SCFM2.SA.XV with NAD at 10 µg/mL or SCFM3. Optical density was measured at 600 nm, and the mean and standard deviation are represented by the solid circle and were calculated over three independent experiments. An ANOVA was run to test the effect of media variation on carrying capacity and growth rate. There was a significant difference in carrying capacity between media types (NCTC 4560 *F*_3,42_ = 8.789, *p* = 1.218*e*^−04^, NCTC 12975 *F*_3,42_ = 82.3191, *p* = *<* 0.001, NCTC 12699 *F*_3,42_ = 21.4965, *p* = *<*0.001) and NCTC 11315 *F*_3,42_ = 650.81, *p* = *<* 0.001) as well as growth rate (NCTC 4560 *F*_3,39_ = 22.494, *p* = *<* 0.001, NCTC 12975 *F*_3,38_ = 11.578, *p* = *<* 0.001, NCTC 12699 *F*_3,42_ = 49.1127, *p* = *<* 0.001 and NCTC 11315 *F*_3,42_ = 15.797, *p* = *<* 0.001). Post-hoc Dunnett analysis identified which media variations resulted in a significant change in K compared to SCFM2.SA.XV. SCFM2.SA.XV without eDNA significantly reduced K in all strains (all *p <* 0.001). There was an increase in K for NCTC 11315 when grown in SCFM3 (*p <* 0.001) and SCFM2.SA.XV with 10 *µ*g/mL NAD (*p <* 0.001). In SCFM3, there was a decrease in K of NCTC 12975 (*p* = 0.034). For r, SCFM2.SA.XV without eDNA significantly reduced r in all strains (all *p <* 0.001). SCFM2.SA.XV with 10 *µ*g/mL NAD reduced the r of NCTC 4560 (*p* = 0.00137).

NAD is an essential growth factor of NTHi, and can reach up to 15 *µ*M in CF sputum, which equates to around 10 *µ*g/mL and as a result is a component of the SCFM3 recipe [47]. Therefore, we also tested whether increasing NAD in SCFM2.SA.XV up to 10 *µ*g/mL, alongside growth in modified SCFM3 which contains purines (adenine, guanine, xanthine and hypoxanthine), NAD+ and p-aminobenzoic acid, would enhance growth of NTHi.

There was a significant increase in carrying capacity for NCTC 11315 when grown in SCFM3 but a decrease for NCTC 12975. Also, 10 *µ*g/mL NAD only enhanced growth of one strain, NCTC 11315, by raising carrying capacity. Increased NAD significantly reduced the growth rate of NCTC 4560. These results suggest while the presence of NAD is important for NTHi growth, it reaches saturating effects at low concentrations.

Overall, these results highlight that SCFM2.SA.XV remains as the most optimal SCFM variant for NTHi growth, and more so, that the eDNA component of SCFM2.SA.XV plays a significant role in supporting the growth of NTHi *in vitro* in SCFM. Although not investigated in this study, eDNA concentrations could be varied to further optimise SCFM2.SA.XV in biofilm studies, as biofilm structure is thought to be highly sensitive to eDNA concentrations (seen in *P. aeruginosa*) [24], and changes in concentration can be the difference between small, but well-developed biofilms, or gelatinous masses and scattered cells. Furthermore, the source of eDNA may influence growth as a result of differences in DNA fragment length incorporated into biofilms.

### Growth of clinical *H. influenzae* isolates was enhanced when grown in SCFM2.SA.XV, demonstrated by increased carrying capacity

The adequacy of laboratory strains to replicate real-world pathogenesis is debated. Many laboratory strains have been sub-cultured over decades since their first isolation, and may have lost important characteristics [69]. We obtained twenty-four clinical NTHi isolates from The Swiss Cystic Fibrosis Infant Lung Development (SCILD) cohort [96][97], and their growth profiles were investigated over 24 hours in SCFM2.SA.XV. We hypothesized that if SCFM2.SA.XV replicated an *in vivo* CF lung environment well, that the media should be able to support the growth of clinical isolates.

We found that all isolates grew in SCFM2.SA.XV. Four isolates (ID41, ID42, ID47 and ID55) had a significantly higher carrying capacity than NCTC 4560, whilst three isolates (ID8, ID21 and ID37) isolates had a significantly lower carrying capacity (**Figure 5A**).

**Figure 5:**
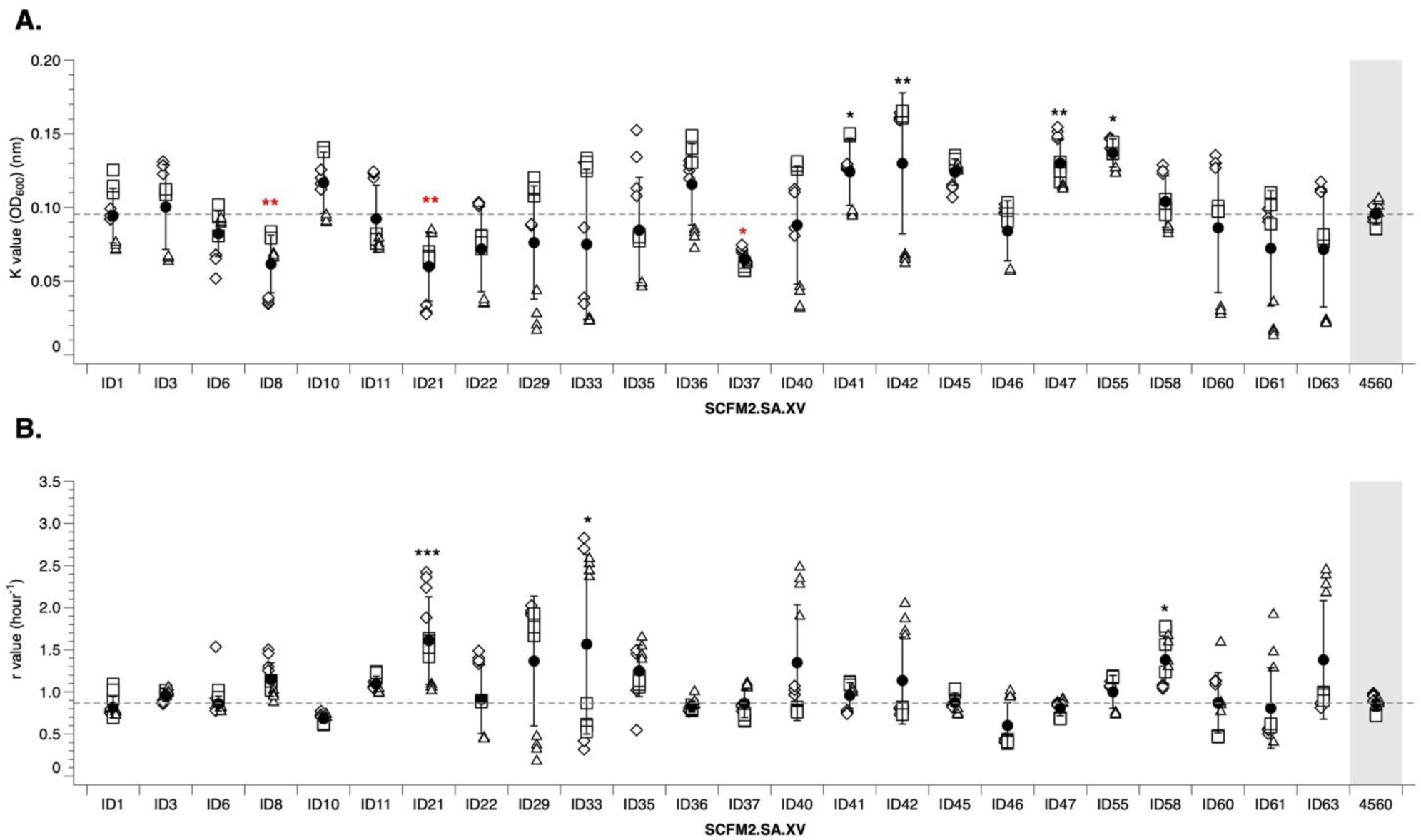
The influence of SCFM2.SA.XV on the growth profiles of clinical *H. influenzae* isolates. NTHi CF clinical isolates were grown statically for 24 hours in SCFM2.SA.XV and both the **A.** carrying capacity and **B.** growth rate were calculated and compared to that of laboratory strain NCTC 4560. Optical density was measured at 600 nm, and the mean and standard deviation are represented by the solid circle and were calculated over three independent experiments. An ANOVA was run to test the effect of isolate on carrying capacity and growth rate (K *F*_24,271_ = 13.376 *p* = *<* 0.001, r *F*_24,271_ = 5.0137 *p* = *<* 0.001). Post-hoc Dunnett analysis identified which clinical isolates had a significant change in K or r compared to NCTC 4560 laboratory strain. 4 clinical isolates had significantly higher carrying capacity than NCTC 4560 which were ID41 (*p* = 0.04597), ID42 (*p* = 0.00692), ID47 (*p* = 0.00724) and ID55 (*p* = *<*0.001), whilst 3 isolates had a significantly lower carrying capacity, which were ID8 (*p* = 0.00321), ID21 (*p* = 0.00692) and ID37 (*p* = 0.0104). 3 isolates had a significantly higher growth rate than NCTC 4560 which were ID21 (*p* = *<*0.001), ID33 (*p* = 0.0311), and ID58 (*p* = 0.0357).

When investigating growth rate, three isolates (ID21, ID33 and ID48) had a significantly higher growth rate than NCTC 4560 in SCFM2.SA.XV (**Figure 5B**). These results suggest that only some NTHi clinical isolates are better adapted to grow in SCFM than a laboratory strain. Growth curves are available in Supplementary Figure 5.

Only isolate ID21 demonstrated both a significantly higher growth rate and lower carrying capacity than NCTC 4560. Potentially, there may be a trade-off such that ID21 is better adapted to utilise SCFM2.SA.XV substrates quickly, but as a result, reaches a lower carrying capacity.

Sequencing of the clinical isolates is required to identify genetic differences between isolates which may result in enhanced growth in SCFM2.SA.XV, and to clarify whether these NTHi isolates are the same strain persisting in one patient, as it is common for individuals to remain colonised with an initial *H. influenzae* strain for up to 2 years without a strain turnover [70].

Overall, it appears carrying capacity of clinical isolates in SCFM2.SA.XV is more variable than growth rate, with larger standard deviation error bars for K than for r.

### Growth of NTHi in *ex vivo* pig lung model with SCFM2.SA.XV

The presence of lung tissue has previously been shown to enhance growth of *S. aureus* and *P. aeruginosa* in an *ex-vivo* pig lung model which utilises pig lungs and SCFM to mimic the CF lung environment [41][42]. However, there is no published study investigating NTHi infection in an EVPL model. In our model, we do not completely sterilise the pig lung tissue prior to use and so endogenous bacteria (pig commensals or environmental bacteria temporarily present in the lungs) can be present. We found that *H. influenzae* is not morphologically very distinguishable from these bacteria by colony morphology. Because there is a lack selective media for *H. influenzae*, a GFP-tagged, streptomycin resistant NTHi, NTHi-375 [51] was used to facilitate identification of NHTi growth in the lung model. To provide a control for this method, a GFP-tagged strain of *P. aeruginosa*, which is known to grow to high density and form biofilm in the EVPL model, was also assayed. An ampicillin + gentamicin wash of the lung tissue was added to previous model preparation methods, to attempt to reduce the titre of endogenous bacteria, before infecting with GFP-tagged NTHi or *P.aeruginosa*, incubating, and plating on VBC sBHI agar (+ 2 µg/mL streptomycin for NTHi). Preliminary work highlighted that this antibiotic wash and plating combination was the most successful for limiting endogenous bacteria growth in the EVPL and on plates used to enumerate target bacteria, respectively. As NTHi is able to invade both bronchioles [71] and alveolar regions of the lung, [15], both bronchiolar and alveolar tissue was dissected from the pig lung and infected to determine whether infection site affected the ability of NTHi to grow.

Our washing step did not completely remove endogenous bacteria, as shown by the high bacterial titre in uninfected tissues (Supplementary Figure 6). Furthermore, endogenous bacteria grew on the selective agar plates, so again we could not reliably distinguish NTHi.

The fluorescence profiles of each tissue section were measured at 0 hours, 24 hours and 48 hours post infection to attempt to visualize NTHi colonization of the lung tissue compared to un-infected tissue (**Figure 6**). Uninfected tissue presented some background fluorescence due to the auto-fluorescence of collagen. However, fluorescence of *NHTi375*-infected tissue did not exceed fluorescence of uninfected tissue at any time point, unlike *P. aeruginosa* for which growth was clearly visible. This suggests that NTHi is unable to establish itself in the EVPL model, despite the presence of optimised SCFM. We hypothesise that this is due to an inability to out-compete endogenous bacteria, but maybe also be because NTHi is an exclusively human pathogen. Further model optimisation is needed to determine whether the EVPL model can be used with NTHi.

**Figure 6:**
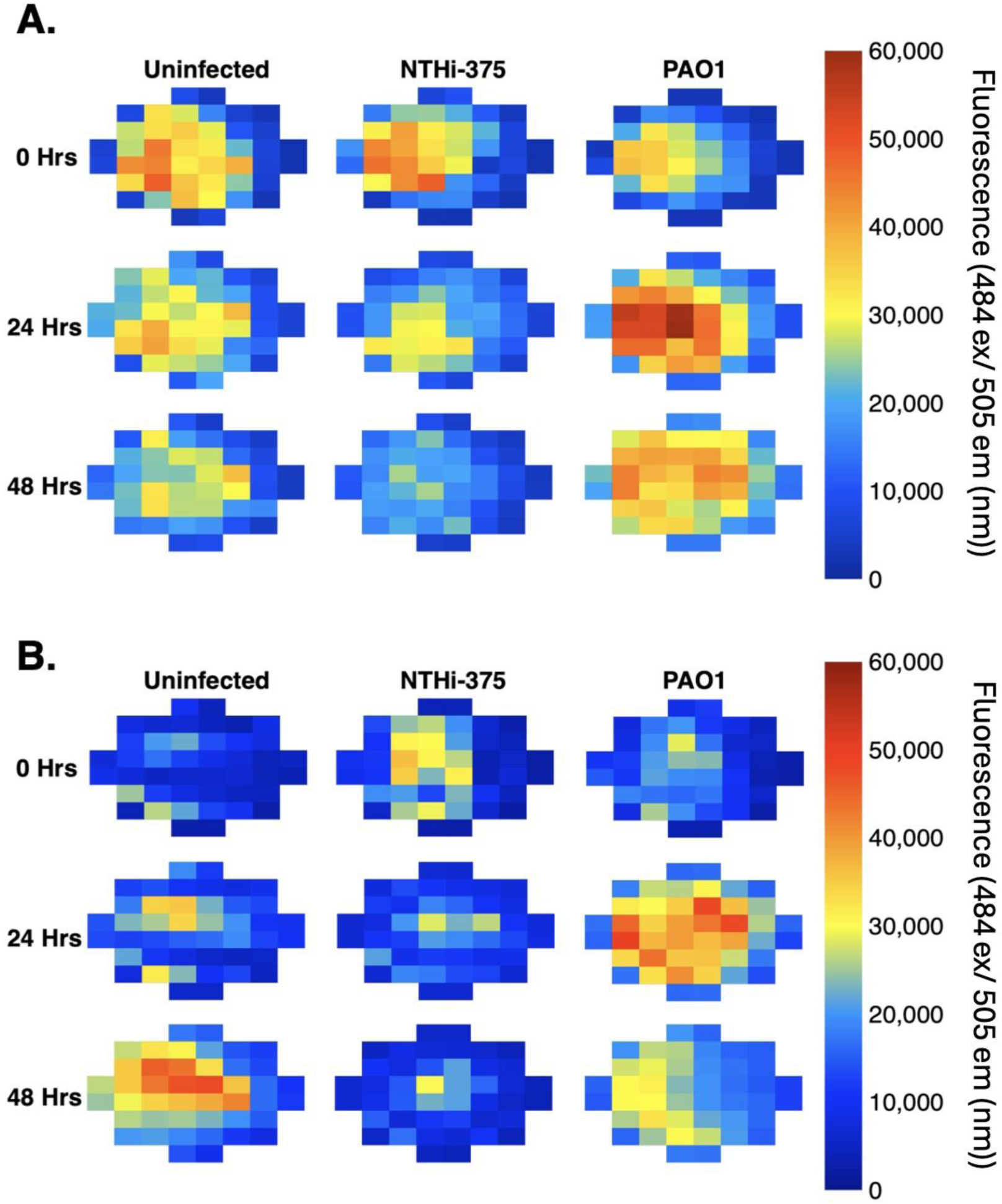
Fluorescence profiles of NTHi-375 in an EVPL model. Bacterial growth was tested on bronchiole lung tissue and alveolar tissue in SCFM2.SA.XV and plating on selective VBC-sBHI agar. Growth of a GFP-tagged PAO1 was used as a positive control. The **A.** fluorescence on bronchiole tissue and **B.** alveolar tissue at 0, 24 hours and 48 hours was monitored. Fluorescence was measured at 484 (nm) ex / 505 (nm) em.

### Lung homogenate media enhances biofilm growth of some NTHi CF isolates over SCFM2.SA.XV alone

Since the presence of lung tissue has shown to be beneficial to the growth of other CF pathogens [42], but the presence of endogenous bacteria seemingly prevented NTHi establishing in the model, lung homogenate media was created by homogenising fresh, uninfected lung tissue in SCFM2.SA.XV and filter sterilising. This retains the presence of host factors, whilst removing potential competitors. We investigated whether lung homogenate media enhanced *in vitro* biofilm formation of both NTHi laboratory isolates and CF clinical isolates through crystal violet staining after 48 hours of static growth. Nine clinical isolates which had previously demonstrated enhanced growth rate or carrying capacity in SCFM2.SA.XV were used for this experiment.

We found that lung homogenate significantly increased biofilm formation in four of the clinical isolates (ID21, ID42, ID55 and ID58), alongside laboratory strain NCTC 4560, when compared to SCFM2.SA.XV alone. Meanwhile, a significant decrease in biofilm formation was seen with ID33. The other isolates were unaffected by the presence of lung homogenate (**Figure 7**).

**Figure 7:**
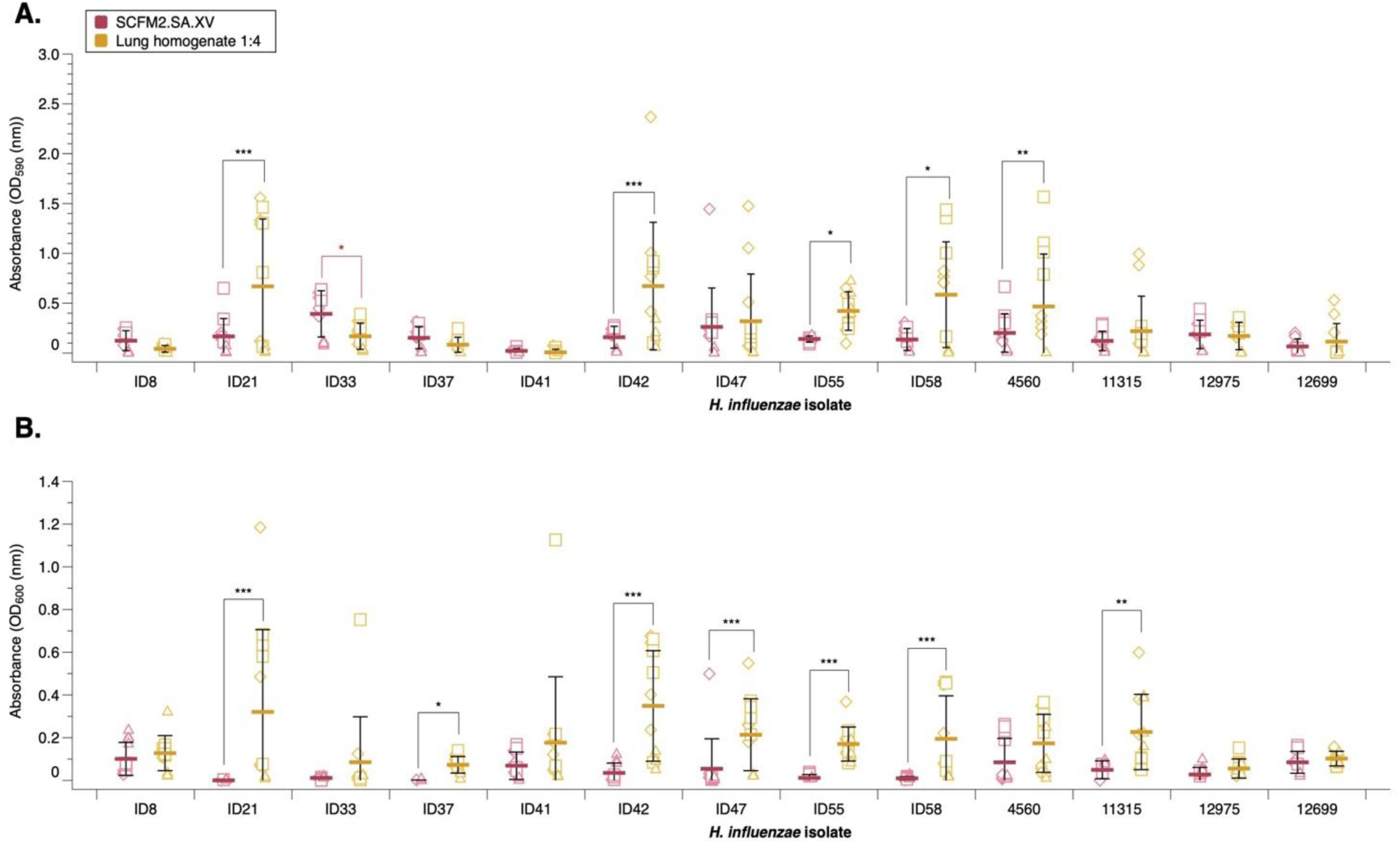
48-hour biofilm and planktonic growth of NTHi laboratory and clinical CF isolates when grown in lung homogenate media. **A**. Biofilm growth and **B**. Planktonic growth were measured by crystal violet staining and OD_600_ respectively. The mean and standard deviation were calculated from 3 independent experiments and represented by the solid line. Significance was defined as *p* ≤ 0.05. An ANOVA was run to test the effect of media type and isolate/strain on biofilm and planktonic growth. There was a significant effect on biofilm formation by media (*F*_1,249_ = 12.0637 *p* = *<* 0.001), strains (*F*_12,249_ = 4.9871 *p* = *<* 0.001) and an interaction strain*media (*F*_12,249_ = 3.0237 *p* = *<* 0.001). There was also a significant change in planktonic growth between media (ANOVA: *F*_1,251_ = 76.1778 *p* = *<* 0.001), strains (ANOVA: *F*_12,251_ = 3.4339 *p* = *<* 0.001) and an interaction strain*media (ANOVA: *F*_12,251_ = 2.9556 *p* = *<* 0.001). Post-hoc analysis using estimated marginal means for pairwise comparisons highlighted which media*strain interactions were significant. There was a significant increase in biofilm formation when grown in lung homogenate media with ID21 (*p* = 0.0007), ID42 (*p* = 0.0009), ID55 (*p* = 0.0262), ID58 (*p* = 0.0158) and NCTC 4560 (*p* = 0.0056). A significant decrease was noted with ID33 (*p* = 0.0348). Planktonic growth was significantly increased in ID21 (*p* = ≤0.0001), ID42 (*p* = ≤0.0001), ID47 (*p* = 0.0002), ID55 (*p* = 0.0005), ID58 (*p* = 0.0001) and NCTC 11315 (*p* = 0.0077).

Lung homogenate media also significantly enhanced planktonic growth of five clinical isolates (ID21, ID42, ID47, ID55 and ID58) and the laboratory strain NCTC 11315. Overall, four of the nine clinical isolates tested (ID21, ID42, ID55 and ID58) had a significant increase in both biofilm and planktonic growth when grown in lung homogenate media compared to SCFM2.SA.XV alone. These results suggest that the presence of lung components is capable of enhancing biofilm formation and planktonic growth of some clinical isolates. However, the growth of replica cultures when in lung homogenate media was much more variable than in SCFM2.SA.XV alone, suggesting the usage of lung homogenate media results in less reproducible growth.

## Discussion

Cystic fibrosis imposes a large economic burden on healthcare systems, individuals with CF and their caregivers. The burden is thought to be underestimated, with many studies only reporting healthcare costs [72]. The increasing prevalence of NTHi [73][74], alongside the changing landscape of CF lung infection with the availability of CFTR modulators, may place a greater importance on handling and treating NTHi infections in people with CF [8][9][10][11] [75].

There are increasing concerns about the relevance of standard nutrient-rich laboratory media to replicate *in vivo* conditions [76] [77] [78]. Host-mimicking media such as SCFM have been demonstrated to better recapitulate the growth dynamics of pathogens such as *P. aeruginosa* and *S. aureus* in the CF lung [38][47] [46]. However, to our knowledge, the growth of the early CF pathogen *H. influenzae* has not previously been studied in any form of CF mimicking media *in vitro*. In this study, we present data on the planktonic growth profiles and biofilm formation of NTHi laboratory and clinical CF isolates in different synthetic sputum media *in vitro*, and present a partially optimised SCFM for usage with NTHi *in vitro* : SCFM2 with mucin replaced by N-acetylneuraminic acid, supplemented with NAD and hemin.

We found that all NTHi grew poorly planktonically in SCFM1. This is the simplest form of SCFM we used, and lacks common CF sputum components such as mucin and eDNA. The carrying capacity and growth rate of NTHi was significantly improved when grown in modified SCFM2 (SCFM2.SA), which contained additional components including sialic acid, GlcNAc and eDNA. We also investigated SCFM2.SA with a source of NAD and hemin either directly, or in the form of FBS. We found that FBS as a source of NAD and hemin hindered biofilm formation, and instead promoted planktonic growth, whilst NAD and hemin as a supplement directly appeared to increase extracellular matrix production, particularly in the laboratory strains NTHi NCTC 4560 and NCTC 11315. Potentially, different ratios of NAD and hemin could be investigated to further optimise SCFM. The negative effect of FBS on biofilms has previously been documented with *P. aeruginosa* [79] as well as *S. aureus* alone and in a polymicrobial biofilm with *Candida albicans* [80]. Components of FBS such as the serum proteins lactoferrin, apo-transferrin, and albumin may have contributed the inhibition of biofilm formation [81] [79] [82].

We found that the presence of eDNA in SCFM2.SA.XV was crucial for the enhanced the growth of NTHi. We found that both the growth rate and carrying capacity of NTHi were significantly reduced when eDNA was omitted from SCFM2.SA.XV. Extracellular DNA has been shown to be present within the matrix of biofilms and is essential for biofilm maintenance of *Haemophilus influenzae* in an otitis media model, and the structure of an otitis media isolate [83][84] [85]. Our results support the idea that eDNA is crucial for NTHi growth and biofilm formation, and the presence of eDNA is also important in the cystic fibrosis lung. The eDNA used as part of SCFM is of low molecular weight (100-200b), but eDNA fragments *in vivo* can vary widely in size: DNA released from cell lysis can be up to several kilobases, while secreted DNA is often smaller. Therefore, we suggest exploring whether alternate sources of eDNA, with varying lengths, also influence the growth of NTHi [99][100].

We also found that replacing bovine maxillary mucin specified in the original SCFM2 recipe with the sialic acid N-acetylneuraminic acid increased the growth rate and carrying capacity of NTHi laboratory strains when compared to SCFM1. NTHi is unable to synthesise its own sialic acid, and therefore depends on uptake of sialic acid from the host [86]. Sialic acid has been shown to be important in the matrix formation of NTHi biofilms through a backbone of polysaccharide and 2,6 sialic acid [87], and NTHi deficient in the ability to sialylate its lipooligosaccharide have been shown to exhibit an impaired ability to form biofilms *in vitro* [88] [87] [37]. Our results align with the idea that biofilm formation of NTHi is enhanced in media containing sialic acid, and that sialic acid is an important component for NTHi biofilm formation [87][29][89][90]. As such, we deemed SCFM2.SA.XV the most optimal media for usage with NTHi as it supports biofilm formation, and *Haemophilus influenzae* isolated from people with CF have been shown to be strong biofilm producers *in vitro* in the literature [91].

We then went on to investigate the ability of NTHi to form biofilms *ex vivo*. We attempted to grow a GFP-tagged isolate, NTHi-375 in an EVPL model, to replicate both the structural and biochemical environment in the CF lung [48] [54]. However, we were unable to successfully grow or identify the presence of NTHi on either bronchiole or alveolar tissue by fluorimetry, and we could not culture NTHi despite the usage of selective agar plates. NTHi is an obligate human pathogen, and does not naturally infect pigs [92]. As such, it is likely this may have affected the ability of NTHi to colonise the lung tissue. We also found that the specificity of NTHi isolation agar was poor. In the literature, haemin-bacitracin chocolate agar is only able to isolate *H. influenzae* from CF sputa at a rate of 13.7%, and cefsulodin-chocolate agar at 17.8% [93]. We found chocolate agar supplemented with vancomycin, bacitracin and clindamycin (VBC) was the most successful at inhibiting endogenous bacteria present on the lung tissue, which aligns with literature results reporting an isolation rate of 59.9% [94]. However, when compared to the specificity of mannitol salt agar for *S. aureus*, which is as high as 97.6%, it is clear there is a lack of selective media to isolate *H. influenzae* from respiratory cultures [95]. We also found that even washing the lung tissue with high concentrations of ampicillin and gentamicin prior to infection did not remove a sufficient level of endogenous bacteria, which may have interfered with NTHi colonisation. Other pathogens including *P. aeruginosa* and *S. aureus* have been successfully grown in the EVPL model [41] [42]. *P. aeruginosa* has been shown to easily out-compete endogenous bacteria present on the bronchiole tissue sections [41], and can be readily isolated using selective agar, whilst *S. aureus* could successfully be distinguished from endogenous bacteria using through plating on highly selective mannitol salt agar [42]. As such, further work is required to better isolate NTHi in the presence of endogenous lung bacteria, and we recommend using an NTHi isolate with an antibiotic resistance marker to improve isolation. However, it is important to note that this may interfere with antibiotic susceptibility studies due to antagonism or synergy between antibiotics used for selection and those under investigation, and resistant mutants may have altered fitness over their wild-type variant.

As the presence of lung tissue has been shown to be important in enhancing the growth of other CF pathogens [42] [41], we added lung homogenate into SCFM2.SA.XV, to keep important components of the lung tissue *in vitro*, whilst removing endogenous bacteria. We found that lung homogenate media enhanced biofilm formation of some clinical NTHi isolates, suggesting that there are components in the lung tissue that contribute to the growth of NTHi and these are not captured in any published SCFM recipe.

We have optimised SCFM in the form of SCFM2.SA.XV for the usage with NTHi, and have highlighted the importance of the presence of eDNA in supporting NTHi growth and biofilm formation. However, further work is required to take NTHi into the EVPL model, including higher specificity isolation agar, and removal of endogenous bacteria. We suggest that proteomic/metabolomic analysis of the lung homogenate media would help suggest a new formulation of SCFM for better replicating the CF lung environment to facilitate further research into NTHi biology.

## Conclusion

An array of synthetic CF sputum media has been developed to attempt to capture the complex nutritional composition of CF sputum [38][46][47]. We found that additions of eDNA, sialic acid and NAD and hemin to SCFM were crucial for NTHi growth and biofilm formation, and that presence of pig lung homogenate enhanced the growth of some clinical NTHi CF isolates even further. Although SCFM2.SA.XV and lung homogenate media improved the growth of NTHi *in vitro*, more work is required to develop a representative CF sputum mimicking media that allows suitable growth of NTHi to better replicate what occurs *in vivo*. We suggest that chemical analysis of the components found in lung homogenate could help to develop an SCFM better suited for studying NTHi microbiology. Our work highlights the need for a representative CF sputum medium that is optimised to support NTHi growth, and we have performed preliminary development of such a medium. Through this development, we can start to better understand the infection dynamics of NTHi, its interactions with other CF pathogens, and its antibiotic susceptibility profile in the CF lung.

## Abbreviations

ANOVA: Analysis of Variance
BHI: brain heart infusion
CF: cystic fibrosis
CFU: colony forming units
CFTR: cystic fibrosis transmembrane conductance regulator
DMSO: dimethylsulfoxide
DNA: deoxyribonucleic acid
DOPC: dioleoylphosphatidylcholine
eDNA: extracellular deoxyribonucleic acid
EVPL: *ex-vivo* pig lung
FBS: fetal bovine serum
GFP: green fluorescent protein
GlcNAc: N-acetyl glucosamine
(HSD): Honestly Significant Difference Honestly significant difference
NAD: nicotinamide adenine dinucleotide
NTHi: Non-typeable *Haemophilus influenzae*
PBS: phosphate buffered saline
SA: sialic acid
sBHI: supplemented brain heart infusion
SCFM: synthetic cystic fibrosis sputum media
UV: ultraviolet
VBC: vancomycin bacitracin clindamycin
XV: hemin and NAD.

## Author contributions

Phoebe Do Carmo Silva: investigation, formal analysis, data curation, writing and editing of original draft. Darryl Hill: conceptualisation, supervision, review and editing. Freya Harrison: conceptualisation, supervision, review and editing.

## Acknowledgements

We thank Karl Staples, Markus Hilty and Ainelen Piazza for supplying bacteria, and Rolf Kümmerli, Lukas Schwyter, Markus Hilty and Karl Staples for helpful discussion of NTHi in CF and feedback on drafts of this work. We also thank Ebba Biotech for the free sample of Ebba BioLight 680 used in this study. The authors would also like to acknowledge the help of the Media Preparation Facility in the School of Life Sciences at the University of Warwick, with special thanks to Cerith Harries and Caroline Stewart.

## Ethical statement

The authors assert that the conducted work involved no experimentation on humans or animals.

