## Supplementary Figure 1 for "Optimising synthetic cystic fibrosis sputum media for growth of non-typeable *Haemophilus influenzae*"

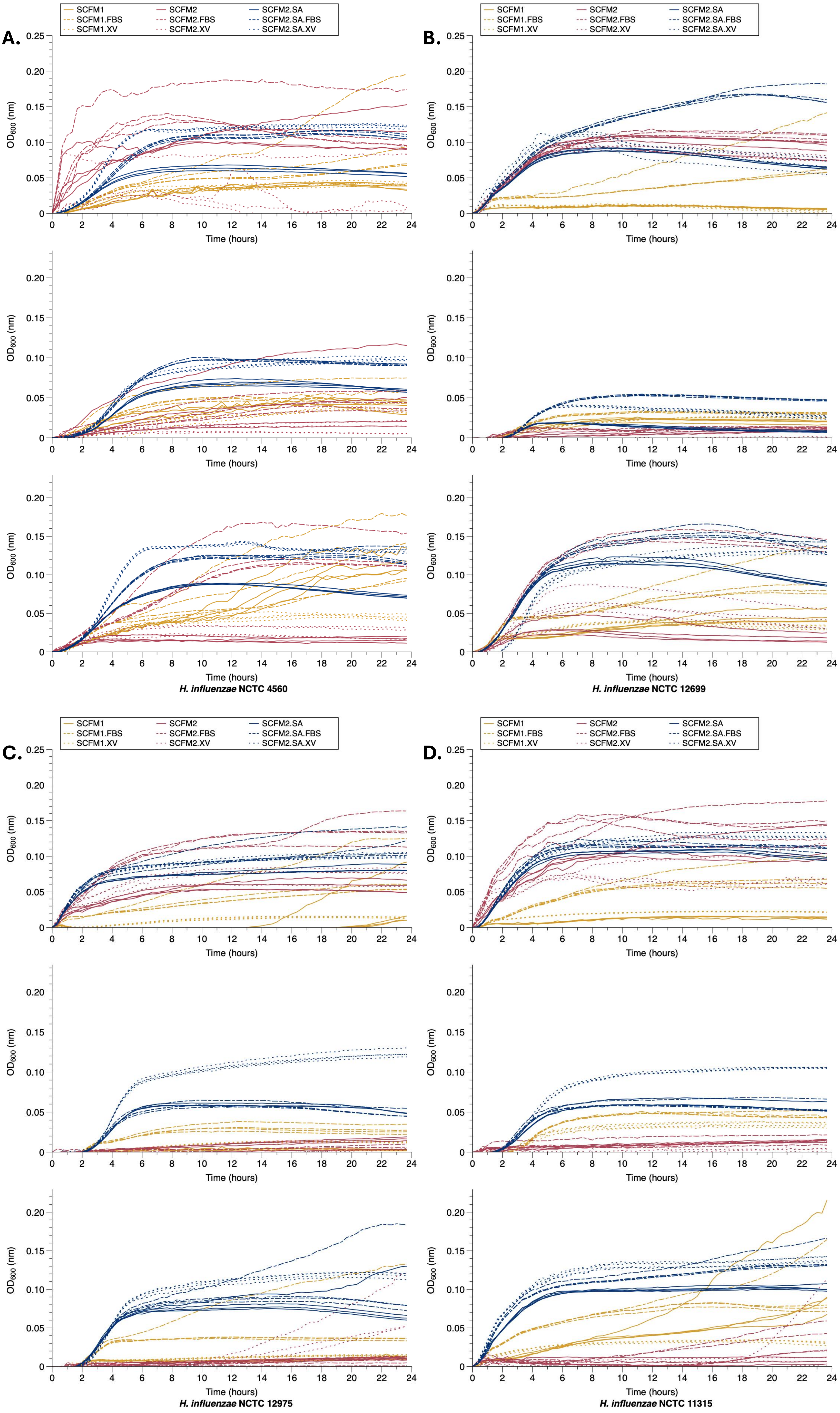


**Supplementary Figure 1 : Growth of *H. influenzae* laboratory strains in variants of SCFM over 24 hours.** Three independent repeats were performed for each lab strain, **A.** *H. influenzae* NCTC 4560, **B.** *H. influenzae* NCTC 12699, **C.** *H. influenzae* NCTC 12975 and **D.** *H. influenzae* NCTC 11315 in 9 variants of media, SCFM1, SCFM2 and modified SCFM2 with sialic acid, alone or supplemented with either FBS or NAD and hemin.
