## Supplementary Figure 2 for "Optimising synthetic cystic fibrosis sputum media for growth of non-typeable *Haemophilus influenzae*"

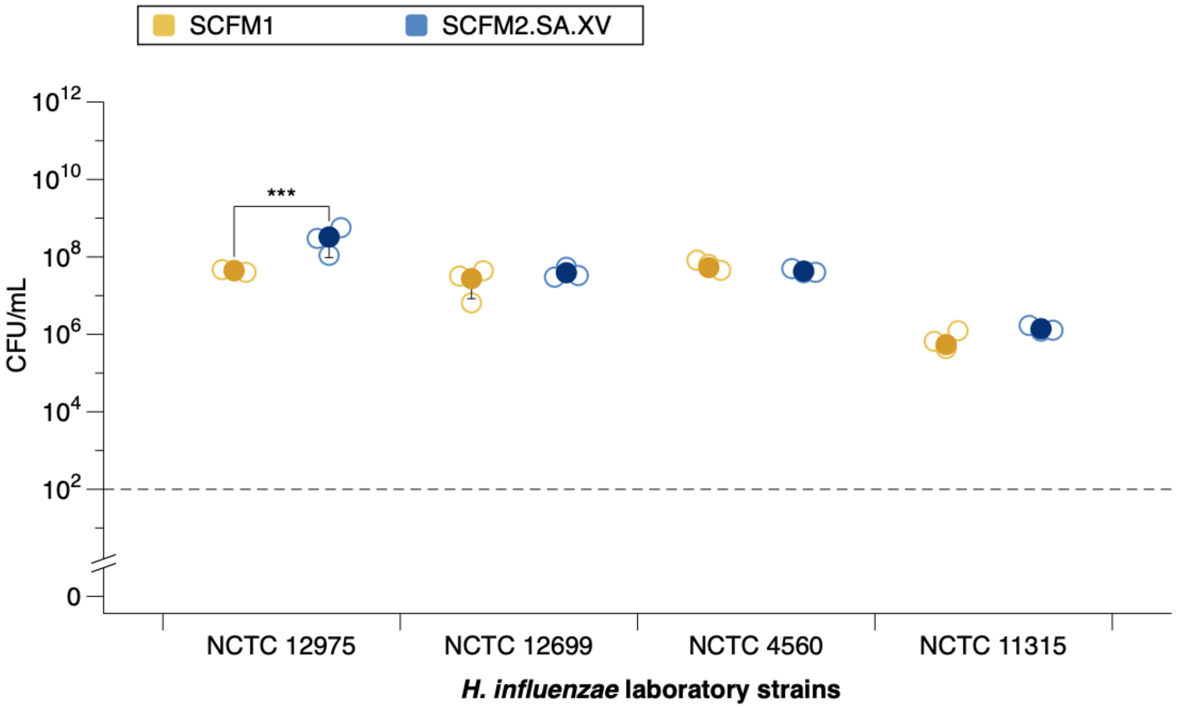


**Supplementary Figure 2: CFU/mL of four laboratory *H. influenzae* strains following 24- hour growth in SCFM1 or SCFM2.SA.XV.**

Mean and standard deviation was calculated from 3 replicates and shown by the solid circle. Significance was defined as $p \leq 0.05$. Limit of detection is represented by the dotted line. There was a statistically significant difference in CFU between between media types (ANOVA: $F_{(1,16)} = 9.57$, $p = 0.007$), strains $F_{(3,16)} = 90.9136$, $p = <0.001$), and an interaction between media and strain $F_{(3,16)} = 4.1696$, $p = 0.0232$). Post-hoc analysis using estimated marginal means for pairwise comparisons highlighted which media*strain interactions were significant. There was an in the CFU of NCTC 12975 when grown in SCFM2.SA.XV compared to SCFM1 ($p = 0.0008$). The CFU of NCTC 11315 was significantly lower in both SCFM1 and SCFM2.SA.XV compared to the CFU of other three laboratory strains grown in these medias (all $p = <0.0001$). NCTC 12975 had a significantly higher CFU when grown in SCFM2.SA.XV (but not SCFM1) compared to NCTC 4560 ($p = 0.0034$), NCTC 12699 ($p = 0.0019$) and NCTC 11315 ($p = <0.0001$).
