## Supplementary Figure 3 for "Optimising synthetic cystic fibrosis sputum media for growth of non-typeable *Haemophilus influenzae*"

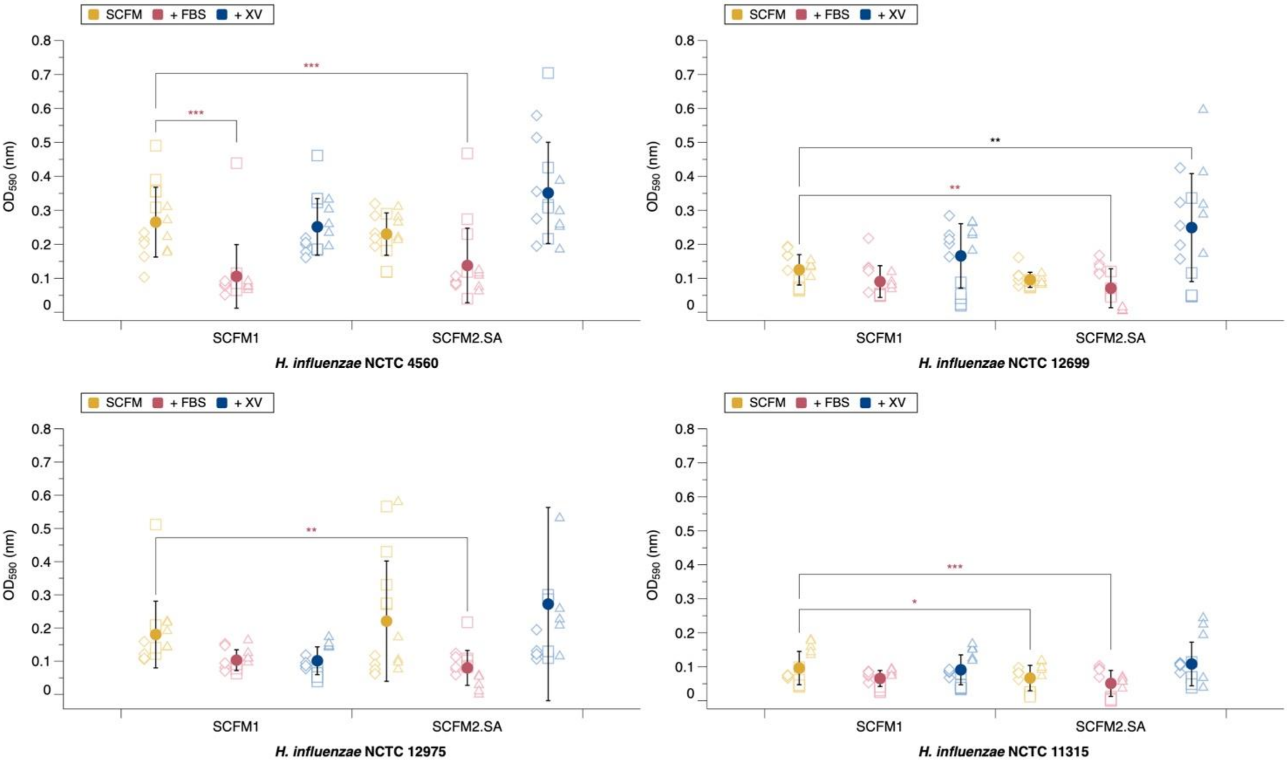


**Supplementary Figure 3: 48-hour biofilm growth of four laboratory *H. influenzae* strains in varying SCFM conditions.** Absorbance was measured at 590 nm. Mean and standard deviation was calculated from 3 independent experiments (demonstrated by different shapes) and shown by the solid circle. Significance was defined as $p \leq 0.05$. An ANOVA was run to test the effect of media variation on biofilm formation for each strain (NCTC 12975 $F_{5,82} = 6.78$, $p = < 0.001$, NCTC 12699 $F_{5,82} = 11.47$, $p = < 0.001$, NCTC 11315 $F_{5,82} = 10.21$, $p = < 0.001$, NCTC 4560 $F_{5,82} = 17.47$, $p < 0.001$). Post-hoc Dunnett analysis identified which media and strain comparisons resulted in a significant change in biofilm formation compared to SCFM1. A significant increase in biofilm growth was seen in NCTC 12699 ($p = 0.002$) when grown in SCFM2.SA.XV. Biofilm formation was significantly reduced in all strains when grown in SCFM2.SA.FBS (NCTC 12975 $p = 0.007$, NCTC 12699 $p = 0.008$, NCTC 11315 $p < 0.001$, NCTC 4560 $p < 0.001$), and a significant decrease was also seen in NCTC 4560 when grown in SCFM1.FBS ($p < 0.001$) and NCTC 11315 when grown in SCFM2.SA ($p = 0.027$).
