## Supplementary Figure 4 for "Optimising synthetic cystic fibrosis sputum media for growth of non-typeable *Haemophilus influenzae*"

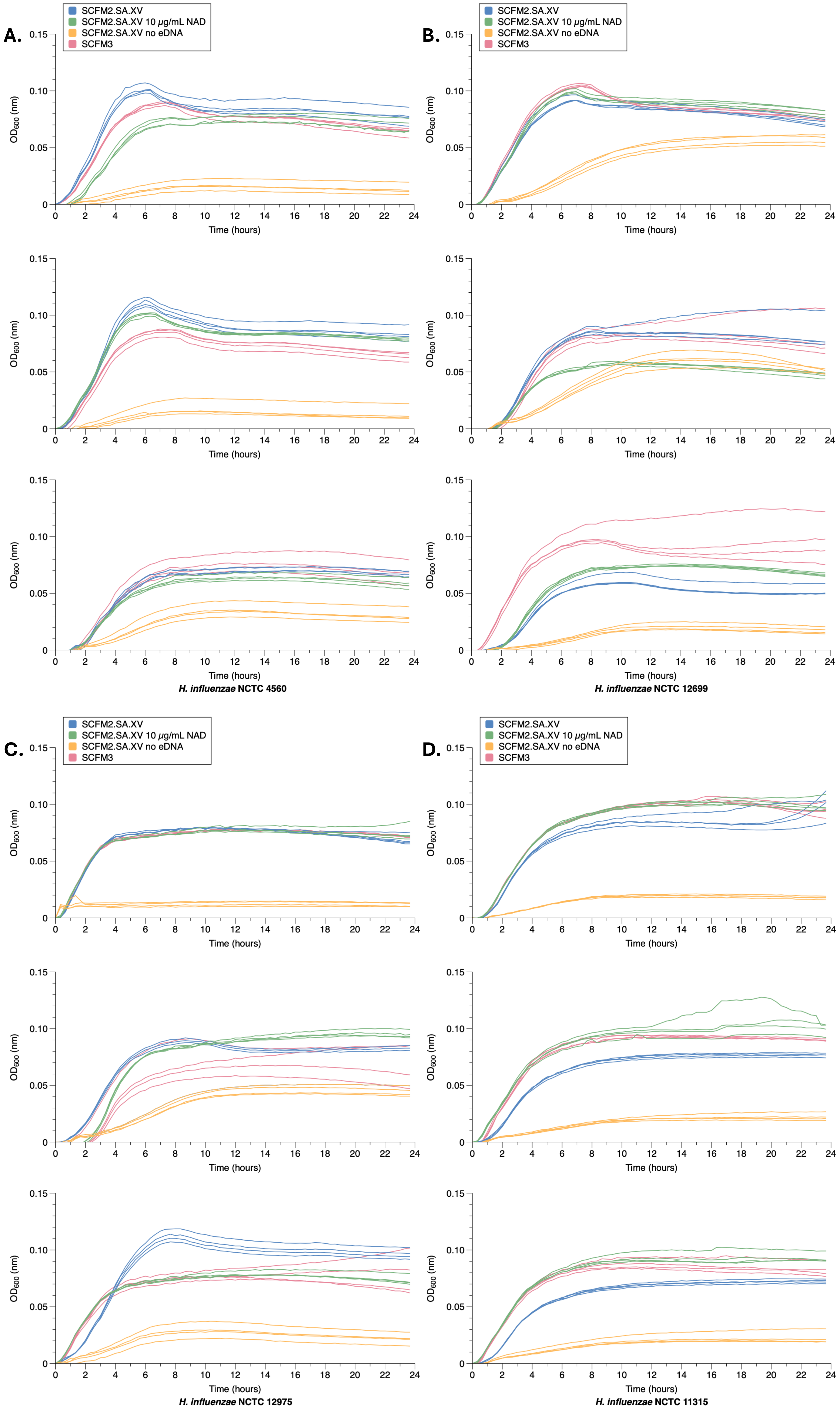


**Supplementary Figure 4: Growth of *H. influenzae* laboratory strains in modifications of SCFM2.SA.XV over 24 hours.** Three independent repeats were performed for each lab strain, **A.** *H. influenzae* NCTC 4560, **B.** *H. influenzae* NCTC 12699, **C.** *H. influenzae* NCTC 12975 and **D.** *H. influenzae* NCTC 11315 in 4 variants of media, SCFM2.SA.XV, SCFM2.SA.XV with 10 µg/mL of NAD, SCFM2.SA.XV with no eDNA and SCFM3.
