## Supplementary Figure 5 for "Optimising synthetic cystic fibrosis sputum media for growth of non-typeable *Haemophilus influenzae*"

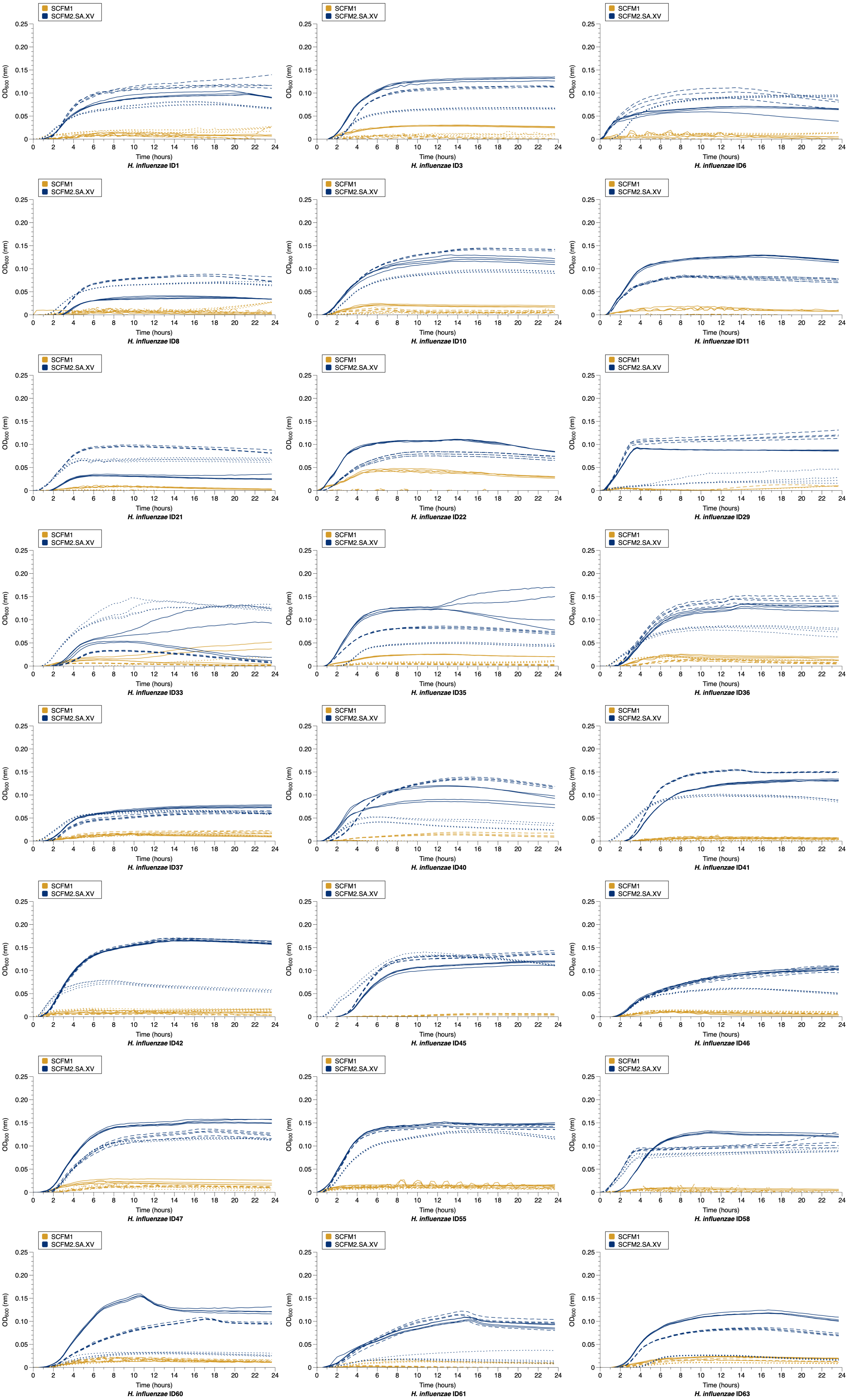


**Supplementary Figure 5: Growth of *H. influenzae* cystic fibrosis clinical isolates in SCFM2.SA.XV and SCFM1 over 24 hours.** Three independent repeats were performed for clinical isolates, with independent repeats shown by the solid line, dashed line and dotted line.
