## Supplementary Figure 6 for "Optimising synthetic cystic fibrosis sputum media for growth of non-typeable *Haemophilus influenzae*"

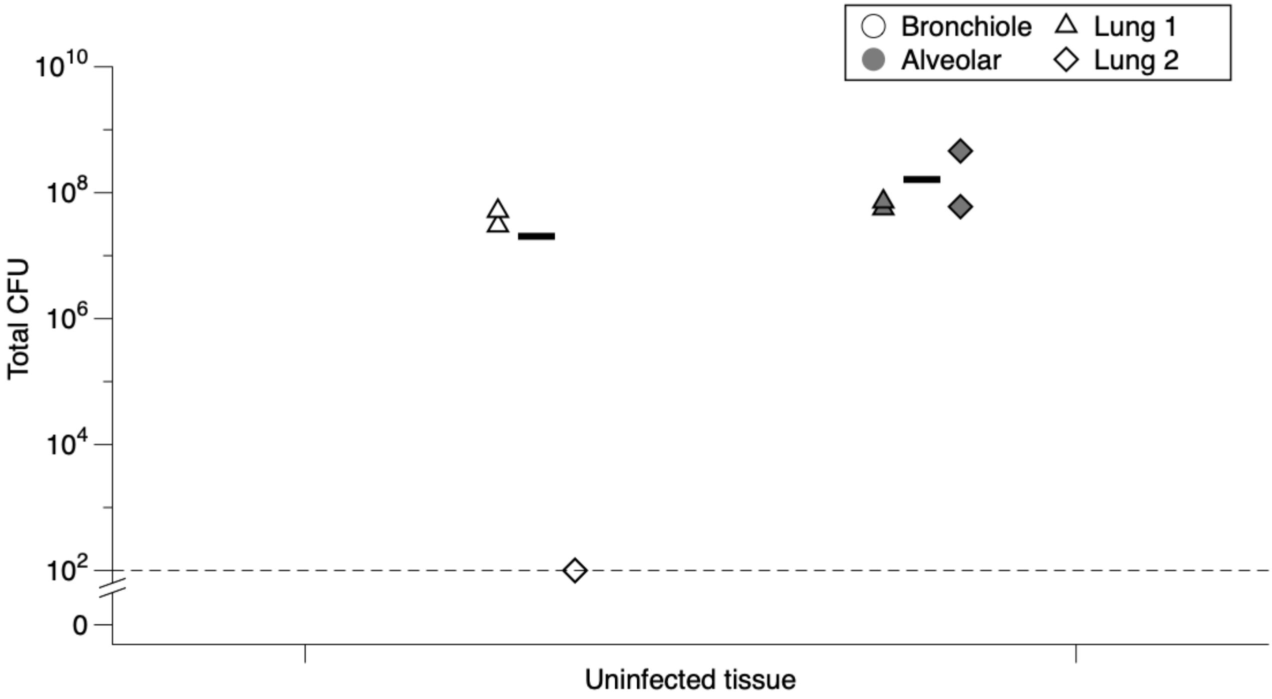


**Supplementary Figure 6: Total CFU of endogenous bacteria on bronchiole and alveolar lung tissue.** Means were calculated from 2 independent experiments (demonstrated by different shapes) and shown by the blue solid line.
